# Long-term evolution of antibiotic persistence in *P. aeruginosa* lung infections

**DOI:** 10.1101/2021.10.14.464434

**Authors:** Melanie Ghoul, Sandra B. Andersen, Helle Krogh Johansen, Lars Jelsbak, Søren Molin, Gabriel Perron, Ashleigh S. Griffin

## Abstract

Pathogenic bacteria respond to antibiotic pressure with the evolution of resistance but survival can also depend on their ability to tolerate antibiotic treatment, known as persistence. While a variety of resistance mechanisms and underlying genetics are well characterised *in vitro* and *in vivo*, the evolution of persistence, and how it interacts with resistance *in situ* is less well understood. We assayed for persistence and resistance with three clinically relevant antibiotics: meropenem, ciprofloxacin and tobramycin, in isolates of *Pseudomonas aeruginosa* from chronic cystic fibrosis lung infections spanning up to forty years of evolution. We find evidence that persistence is under positive selection in the lung and that it can particularly act as an evolutionary stepping stone to resistance. However, this pattern is not universal and depends on the bacterial clone type and antibiotic used, indicating an important role for antibiotic mode of action.

## Introduction

Bacterial persistence is defined as the ability of cells to adopt a phenotype that tolerate and survive exposure to a bactericidal drug concentration (1-3). It contributes to recalcitrant chronic infections even when infecting strains are susceptible to antibiotic treatments (1-3). We must better understand how bacteria respond and adapt to drugs *in situ*, to be able to use them effectively. Most work has focused on resistance, but while a level of consensus has been reached on how to define and quantify persistence (1), the extent to which it is under positive selection in infections from antibiotic treatment and how persistence co-evolves with resistance is still not well understood (4). To characterise long-term evolution of resistance and persistence in infection, we studied isolates of *Pseudomonas aeruginosa* from chronic lung infections of individuals with cystic fibrosis (CF). This infection system is ideal for detecting selection on persistence because long-term sampling has occurred both of lineages of transmissible strains (infecting for 40 years) and within single infections (up to 7 years)(5). The collection of transmissible strains in particular allows us to look at evolutionary history over decades.

In contrast to the antibiotic resistance phenotype, there is little evidence for a clear genetic basis or the molecular mechanisms involved for the persistence trait. A specific gene responsible for persistence has been found only for *Escherichia coli*, while a range of regulatory and metabolic genes and mechanisms that contribute to or correlate with the persistence phenotype have been identified in *Pseudomonas aeruginosa*, as well as other pathogenic species (1, 4, 6-10). In addition, persistent pathogens are often attributed to dormant or slowly dividing cells (11, 12) including those in biofilms (13), although dormancy has been shown to not always be necessary (14). In fact, multiple persister subpopulations that have been induced by different mechanisms may coexist (1). Whether the genetics and mechanisms of persistence are yet unknown or multiple are not required for testing whether this phenotype is under selection, and should not hinder achieving an understanding of how persistence evolves.

Adaptation to antibiotic treatment depends on the frequency of exposure and the encountered concentration of the drug (15, 16). One hypothesis is that persistence is favoured as a bet-hedging strategy (17) when antibiotic concentration is high but exposure infrequent (16, 18-20), and it may evolve in response to intermediate concentrations and facilitate subsequent evolution of resistance (21-24). There is rarely more than 1% of an *in vitro* population in a persistent state at any given time, but this is sufficient to re-colonise once antibiotic pressure has been released (1). If persistence is adaptive we expect to see its frequency increase under strong selection from antibiotics. Could such an increase contribute to failure to treat chronic infections? Persistence in *P. aeruginosa* pathogens was observed repeatedly and early in infection (5, 25) and was found to be selected for prior to resistance *in vitro* (10, 25, 26). This suggests that it is a heritable trait but do we observe similar patterns of its emergence *in situ*?

Here we test how *P. aeruginosa* antibiotic response strategies evolve during infection, by measuring and comparing changes in persistence and resistance to three antibiotics used in the clinic. We use two collections of isolates from individuals with cystic fibrosis (CF), to measure how these traits evolve within individuals and secondly, within lineages of strains transmitting between individuals over extended periods of time. With the first collection, consisting of 97 isolates, we compare environmental isolates that have not been exposed to antibiotics to clinical isolates from early and chronic infections, from 20 individuals. Using another collection of two transmissible strains, we track the evolution of resistance and persistence in 107 clinical isolates sampled longitudinally from 27 individuals over a period of four decades. These comparisons allow us to detect evidence of a role for persistence in the transition from the environment, to acute, and finally chronic infection.

To map the evolution of resistance and persistence during infection we challenge the isolates with three antibiotics used in the clinic, that have different modes of action: ciprofloxacin, a fluoroquinolone that interfere with DNA replication; meropenem, a beta-lactam that inhibit cell-wall synthesis; and tobramycin, an aminoglycoside that inhibit protein synthesis. We further investigate if bacterial survival is specific to the type of antibiotic, or a generalist response correlated across types, and whether there is an interaction between resistance and persistence which may have clinical implications for developing treatment of *P. aeruginosa* lung infections of cystic fibrosis individuals.

## Methods

### Isolate collection

We use two collections of isolates from Danish cystic fibrosis individuals sampled at Rigshospitalet, Copenhagen. One collection covers the first seven years of infection more extensively (27) with the first isolates sampled from twenty individuals (n = 25 acute isolates, 17 clone types) and from three months to seven years into infection (n = 20 chronic isolates, 14 clone types, table S1). Additionally, we include 26 non-lung strains consisting of various lab PAO1 strains (n = 7, phenotypes averaged into one due to genomic similarity), a PA14 isolate and environmentally sampled strains (n = 18, table S1, (28)). We collectively refer to these as non-lung isolates and compare them to the acute and chronic cystic fibrosis isolates.

Another collection consists of transmissible isolates from 27 individuals infected with two clone types that were transmitted within the clinic (57 DK1 isolates from 13 individuals sampled between 1973 and 2012, and 50 DK2 isolates from 18 individuals, sampled between 1972 and 2007; (29-33); see table S2). The DK2 phylogeny suggests that a successful line evolved early, which was transmitted between all subsequent individuals, whereas the DK1 phylogeny is less well resolved with more distinct subclades (Fig. S1A & B). Length of infection for DK1 and DK2 represents time since the first isolate of the clone type was recorded in 1972 or 1973, respectively.

### Quantifying antibiotic resistance

To prepare cultures, we grew freezer stock overnight in liquid LB media at 37 °C with shaking and then standardised cultures to an optical density 0.5 McFarland standard. We then assayed antibiotic resistance in these cultures by swabbing bacterial suspensions onto Mueller-Hinton agar plates with three ETEST plastic strips placed on top in accordance with the manufacturer’s instructions (Liofilchem, Italy). ETEST strips have a predefined gradient of antibiotic concentration (34), which allowed us to measure minimum inhibitory concentration (MIC) values. The concentrations measured were: ciprofloxacin: 0.002-32 µg/ml, meropenem: 0.002-32µg/ml, and tobramycin: 0.016-256µg/ml. We scored the isolates as sensitive, intermediate or resistant according to the species-specific MIC clinical breaking points user guide available from Rosco Diagnostica (http://pishrotashkhis.com/wp-content/uploads/2017/07/Neo-SENSITAB-CLSI-EUCAST-Potency.pdf), and as defined by Clinical and Laboratory Standards Institute (CLSI.org). The MICs were measured on agar plates using strips, which has been shown to be comparable to values obtained in liquid culture (35) and therefore supporting our persistence assays done in liquid media.

### Quantifying persistence

We cultured isolates overnight in liquid LB media and incubated them at 37 °C with shaking. We then standardised the culture to an optical density (OD) of A_600_ = 0.1. From this, we inoculated 2 µl into 200 µl of LB media in 96 well plates and incubated them at 37 °C for 24 hours. As a proxy for growth rate and population size we calculated the mean OD of the replicates after 24 hours of growth in media. After the 24 hours we added antibiotic to each well at a concentration of 100 µg/ml (meropenem, tobramycin, or ciprofloxacin). We incubated the cultures for 24 hours at 37 °C and subsequently plated them out on LB agar plates (see below). Isolates were replicated 8 times for each antibiotic. In the meropenem treatment four isolates that have MIC values above 100 µg/ml were removed from the persister analysis (these isolates were measured using Estrips with a wider range of concentration values reaching a maximum of 256 µg/ml). 35 isolates showed no inhibition zones in the concentration range of the meropenem Estrips we used (0.002-32µg/ml), indicating that their resistance values are above 32 µg/ml. However, we assume that for these isolates, resistance is still below 100 µg/ml (at least 3 times the maximum concentration measured).

Following incubation with antibiotics, we diluted the *P. aeruginosa* cultures 10 and 100-fold using 9 % NaCl solution. Undiluted, 10 and 100-fold dilutions were plated out onto LB agar plates in 5µl drops to count the viable colony forming units (CFUs). Maximum number of CFUs counted was set to 150. For each isolate we calculated the mean CFU and standard deviation around the mean from the replicates at each dilution.

### Persister variation between non-clinical and clinical isolates

We used 1-way ANOVAs to compare levels of persistence in response to antibiotic treatments of the isolates categorised as non-lung, acute and chronic.

### Longitudinal changes in antibiotic persistence and resistance during infection

We analysed the evolution of persistence and resistance for the transmissible clone types DK1 and DK2, using generalized linear models (GLMs), to test the relationship between persistence (as CFU counts) and resistance (as MIC values), with length of infection and initial density after 24h growth.

### Classifying resistance and persistence

The isolates’ MIC for each antibiotic range from 0.008 – 32 µg/ml for ciprofloxacin; 0.006 – 32 µg/ml for meropenem (with the exception of four which have been removed from the persister analysis), and 0.064 -32 µg/ml for tobramycin. We classified the isolates based on the clinically relevant cut-offs (as defined by Clinical and Laboratory Standards Institute, CLSI.org), and with a conservative approach, grouped the intermediately resistant isolates with the susceptible in subsequent analyses. The antibiotic concentration we used for the persister assays was significantly above the MIC, i.e. *ca*. between three to 17 thousand times higher. Any growth (observed as CFU counts) from antibiotic treated cultures are considered to be a result of persistence, defined here as ability to survive exposure to the antibiotic at values above the clinical MIC cut off.

For the analyses we chose to use CFU counts from the 10-fold dilution for all antibiotics to facilitate comparison across antibiotics and optimise the number of isolates with countable CFUs (>1 and <150), without loss of data due to dilution (Table S2). From the tobramycin treatment the plating of undiluted culture yielded most isolates with CFUs, while 10-fold dilution worked best for the ciprofloxacin and meropenem treatments. CFU count is expected to decrease with dilution, however, this also dilutes the antibiotic concentration in the media and we sometimes observe more CFUs at higher dilutions due to cells reviving. We set 75 CFUs as a cut-off, where isolates with counts above this are categorised as “high persisters”, and those below categorised as “low persisters”. There was a bell-shaped relationship between mean CFU count and the value of the standard deviation around the mean, which reflects that isolates with mean CFU counts between 25 and 100 have a large variation between replicates, rather than an intermediate number of CFUs (Fig. S2). The midpoint count of 75 CFUs is at, or above the peak of the standard deviation. Isolates with < 75 CFUs will have more replicates with no CFUs than high CFUs, whereas isolates with >75 CFUs will have more replicates with max CFU.

### Correlation of persistence phenotypes between antibiotics

We want to test whether there is a coupling between changes in persistence and resistance between antibiotic treatments, that is if we e.g. see a change from low to high persistence under ciprofloxacin treatment, does this lead to a matching persistence phenotype observed with meropenem? As an example, in the phylogeny Fig. S3E branch *QE* (marked as “DK2.1”) we see a change in persistence under meropenem, which leads to the persistence phenotypes observed from ciprofloxacin and meropenem treatments being unmatched. On branch *LF* (marked as “DK2.2”) there is a change in persistence under ciprofloxacin treatment so that persistence phenotypes again are matched. In the analysis this is described as two events, where the first results in unmatched persistence phenotypes, the second to matched phenotypes. For all pairwise combinations of antibiotics (ciprofloxacin and meropenem; ciprofloxacin and tobramycin; meropenem and tobramycin) we tallied the changes in persistence (from matched to unmatched or vice versa) across the phylogenies for DK1 and DK2 (Table S3). We calculate the probability of finding the distribution of matched and unmatched events as [P(X ≥ events _observed_) ∼ pois(X; events _expected_) < 0.05], where the expected number of events were half of the total events.

### Transitions between levels of persistence and resistance

To determine if resistance and persistence evolves in a specific order we use the *in vitro* antibiotic treatment data to classify the antibiotic response strategy of each isolate into one of four phenotypic categories based on the CFU counts and MIC values: low persistence | susceptible (LS); low persistence | resistant (LR); high persistence | susceptible (HS) or high persistence | resistant (HP). We mapped the phenotypes of all isolates onto the phylogenies of DK1 and DK2 to identify independent transitions between antibiotic responses. Resistance and persistence evolve and are lost repeatedly in both clone types. For each antibiotic, the most parsimonious order of changes was determined and the phylogenies divided into monophyletic clonal lineages. From these we observed the number and order of transitions (e.g. from susceptible | low persistence to susceptible | high persistence, followed by development of resistance). The probability of finding the observed distribution of changes was calculated as [P(X ≥ events _observed_) ∼ pois(X; events _expected_) < 0.05], where the expected number of events were half of the total number of transitions from/to a given category.

Given the four different phenotypic categories of response strategies and transitions between them, we can compare the number of transitions to and from each. We define a “stable” phenotype as one that has few transitions from it, and “unstable” as one that has many transitions from it. To test whether a phenotype was significantly more, or less, stable than expected given random transitions between phenotypes, we calculated the probability of finding the observed number of transitions from a phenotype compared to the expected number, given as a quarter of the total transitions from any phenotype. We analysed all isolates of DK1 and DK2 together to increase the sample size, and for each antibiotic separately and all together (Table S4).

We tested if there was a difference in maximum OD, and sampling time, of resistant or persistent isolates, and an interaction between the two factors, with a 2 -way ANOVA with Tukey post hoc comparisons.

### Statistical analysis

All statistical analyses and graphs were done in R version 3.3.1 (R core team 2018), using packages broom (36), gridextra (37), ggbeeswarm (38), and tidyverse (39), including dplyr (32), ggplot2 (32), readr (32), and forcats (32).

## Results

### Selection on persistence in infection

#### Patterns of persistence evolution in transmissible lineages

We found strong evidence for positive selection on persistence cell formation in both transmissible strains tested. In the transmissible strain DK2, we found positive selection in all three antibiotics treatments. More specifically, we found a significant increase in persister cell counts per isolate over time in response to all three experimental antibiotic treatments (GLMs, p < 0.05; Table S5, Fig. 1). We first observed high persistence (>75 CFUs per isolate) after 19 years of sampling across patients in response to ciprofloxacin, after 7 years in response to meropenem, and after 12 years for tobramycin. Once evolved, high persistence was maintained in at least 86% of clonal lineages to the end of the sampling period (Fig. S3D, E, F & S9) [ciprofloxacin, n= 24/26, meropenem n= 31/36 and tobramycin n=24/26], suggesting that persistence was either fixed in these populations or maintained in the population via selection.

**Fig. 1.**
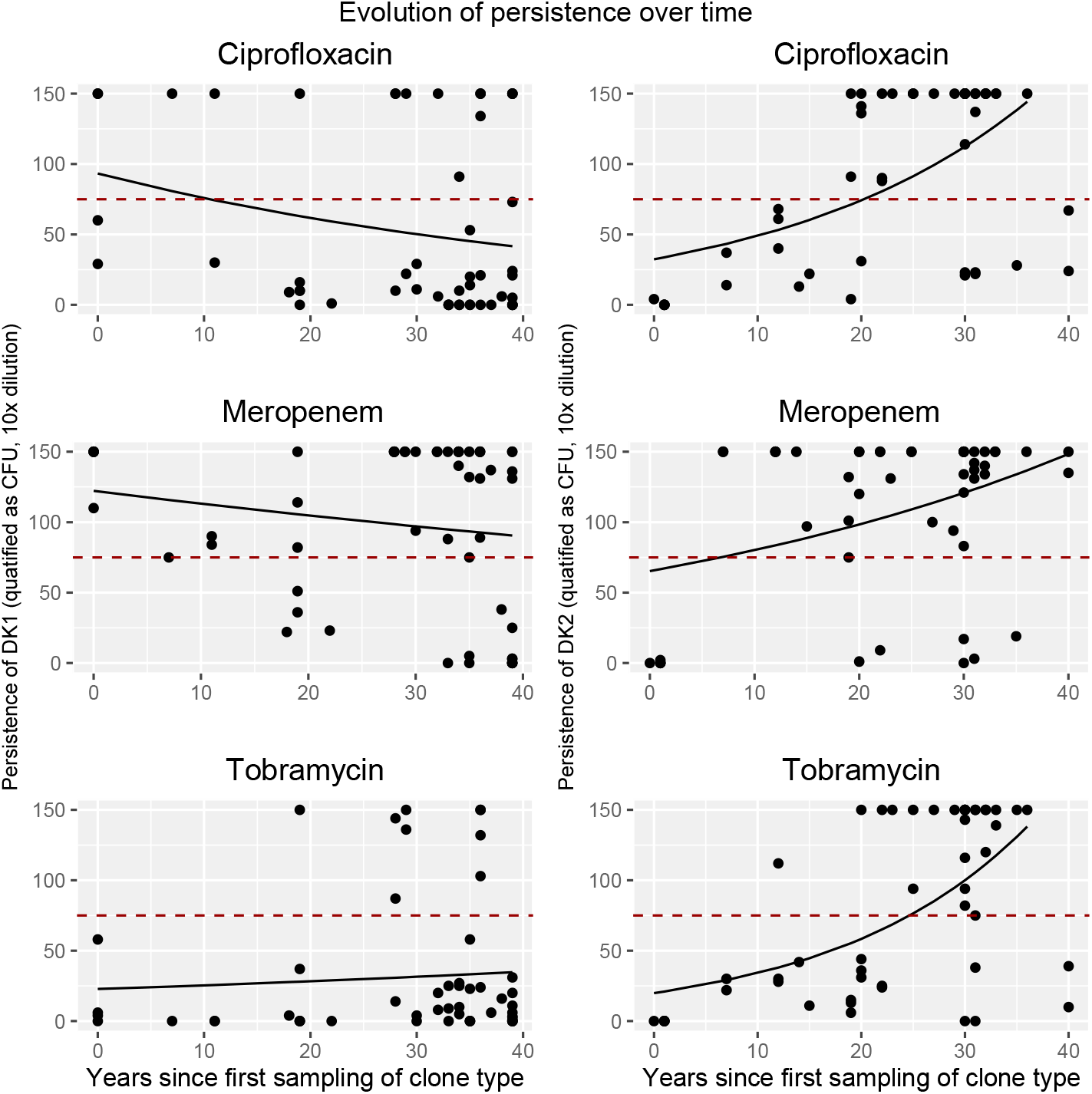
Graphs show number of persister cells per isolate, measured as CFUs, over time for the two different clone types and three different antibiotics, and the GLM fit using time to predict change in CFU. The dashed red line marks the cut-off of 75 CFU to classify isolates as either low or high persisters. On the left, DK1, and on the right, DK2. Persistence was measured as CFU counts at 10 times dilution of culture.

For the DK1 lineage, persister cell production detected in our tobramycin treatment increases significantly with length of infection, with high persistence first observed after 19 years. In contrast, while high persistence is observed in early DK1 isolates in response to ciprofloxacin and meropenem, there is a significant decrease in persistence over time (GLMs, p < 0.05; Table S5, Fig. 1).

#### Persister phenotypes correlate across meropenem and ciprofloxacin in transmissible lineages

After exploring the evolution of persistence to an individual antibiotic we test if there are patterns in isolate response across the three antibiotics. When following clonal lineages on the DK1 and DK2 phylogenies we observe changes in the persistence phenotype going from either low to high or high to low persistence. In pairwise comparisons between antibiotics we find that persister phenotypes in response to meropenem and ciprofloxacin treatments are frequently matched, so that an isolate with high persister under meropenem treatment is also a high persister to ciprofloxacin, or a low persister under both antibiotic treatments. The null model would predict that 50% of transitions would lead to the same persister phenotype and 50% would lead to unmatched persister phenotypes. However, we find that significantly more often than expected by chance, a change in persistence to ciprofloxacin lead to the same persister phenotype as for meropenem, and vice versa [P(X ≥ 27) ∼ pois(X; 14.5) = 0.002]). The tobramycin treatment does not show that changes in the persister phenotype more often than expected by chance leads to a shared phenotype with either meropenem [P(X ≥ 17) ∼ pois(X; 12.5) = 0.13] or ciprofloxacin [P(X ≥ 9) ∼ pois(X; 12.5) = 0.88].

We observe the independent evolution of high persistence to all three antibiotics seven times, and the same for resistance (Fig. S1A & B). For five of these there is an overlap, in that the same isolates show resistance and persistence to all antibiotics.

#### Patterns of persistence evolution at early stages within individual patient infections

Isolates from chronic infections show significantly higher persistence than non-lung isolates detected in the meropenem treatment (F_2,69_ = 15.65, p < 0.001; Fig. 2). We found no significant difference in persistence between classes of isolates under both ciprofloxacin (1-Way ANOVA, F_2,69_ = 0.741, p = 0.48) and tobramycin antibiotic treatment (F_2,69_ = 1.831, p = 0.168, Fig. 2). High persisters were detected in only one non-lung and one acute isolate in response to ciprofloxacin treatment.

**Fig. 2.**
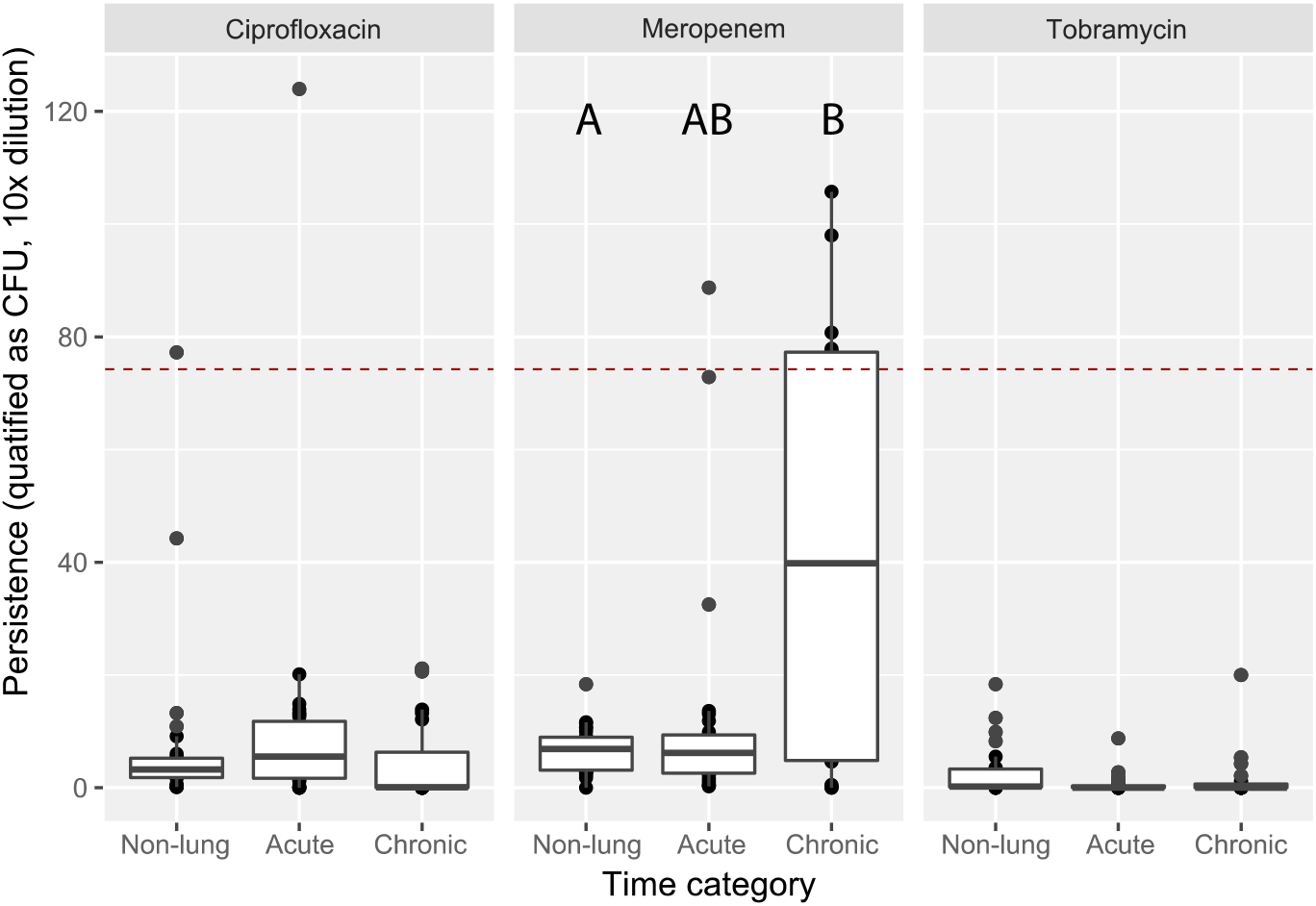
Graphs show persistence of isolates, measured as CFU counts, sampled from three categories in response to three different antibiotics. Boxplots show median CFU count ± 25 percentiles and values as dots. The dashed red line marks the cut-off of 75 CFU to classify isolates as either low or high persisters. Non-lung isolates (n = 26 isolates of 26 clone types); acute isolates, sampled upon first signs of infection which may have ranged within days to a couple weeks (n = 26 isolates of 17 clone types); and chronic isolates, sampled from three months up to seven years after the first detected infection (n = 20 isolates of 14 clone types; see table S1).

### Resistance is under positive selection in infection

For the transmissible clone types, there is a significant increase in antibiotic resistance, measured as MIC, to all three antibiotics, for both clone types analysed together and separately (DK1: 57 isolates sampled between 1973 and 2012; and DK2: 50 isolates sampled between 1972 and 2007; GLMs, p < 0.05; Table S5, Fig. 3, see Fig. S6 for plots split by clone type). However, it takes over a decade for clinically resistant mutants (see methods) to become detectable: resistance to ciprofloxacin is first observed after 20 years of infection, and after 12 years for meropenem and tobramycin. However, it is important to note that no data is available on when treatment with each antibiotic type was initiated in patients, which influences resistance onset.

**Fig. 3:**
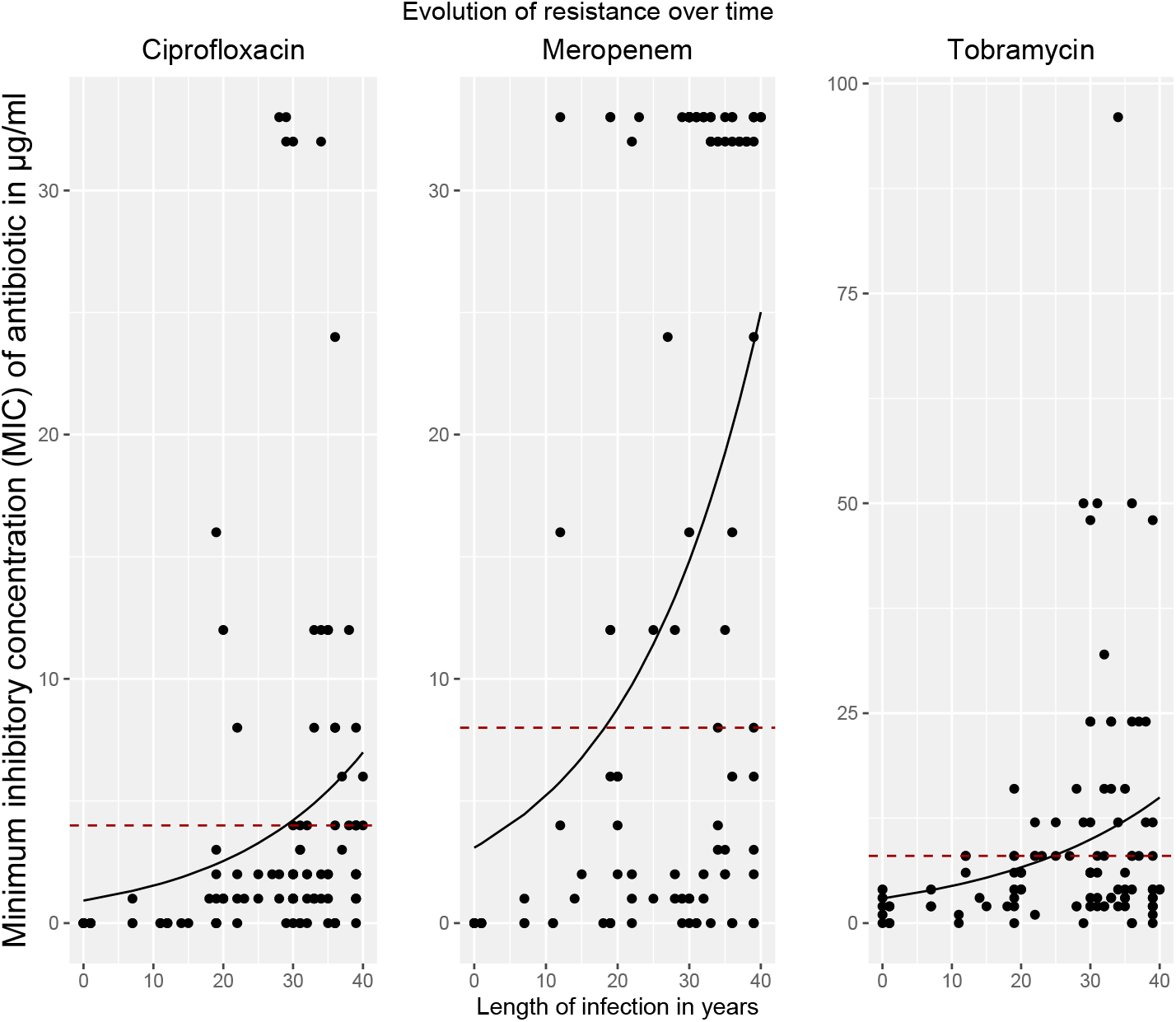
Graphs show change in resistance (MIC) over time in the DK1 and DK2 transmissible clone types combined, to three different antibiotics (GLM fits). Dashed red lines indicate clinical cut-off for classification of resistance (MIC of 4 for ciprofloxacin, and 8 for meropenem and tobramycin). For plots of DK1 and DK2 clone types separately see Fig. S6).

In line with this, all isolates from the non-lung, acute and chronic collection covering the first seven years of infection are clinically susceptible to the three antibiotics, except one of the chronic isolates that was resistant to meropenem (Fig. S4).

### Co-evolution of resistance and persistence within transmissible lineages

#### Selection for persistence precedes resistance

Given that resistance and persistence are under positive selection, we next show how these traits might co-evolve within transmissible lineages. We assessed what the most stable phenotype is, defined as one from which there are fewer than expected transitions to other phenotypes (see methods), and the route taken to get there (Fig.4). The patterns we observe are distinct for the different experimental antibiotic treatments. Meropenem data shows that the resistance | low persistence phenotype is significantly more stable than expected by chance (LR; P(X ≥ 8.25) ∼ pois(X; 1) = 0.002). Resistance in combination with persistence is the stable phenotype under ciprofloxacin treatment (HR; P(X ≥ 7.25) ∼ pois(X; 0) < 0.001). In contrast, the response to tobramycin shows that resistance alone, persistence alone, and the combination of persistence and resistance are all equally stable phenotypes while the susceptible | low persistence phenotype is significantly less stable than expected by chance [LS; P(X ≤ 6.5) ∼ pois(X; 12) = 0.034]; Table 1 & Fig. 4). This is in line with the sampling times of the different categories (Fig. 5).

**Table 1:** Transitions between the different responses to antibiotics. Ciprofloxacin: most transitions are from low persistence | susceptible to high persistence | susceptible, while the stable phenotype (least transited out of) is high persistence | resistant, indicating the role of persistence as a stepping stone to resistance. Meropenem: most transitions occur from high persistence | susceptible to high persistence | resistant, but the least transited from phenotype (stable) is low persistence | resistant. This indicates that resistance is more effective for meropenem survival. Tobramycin: high persistence, without resistance, may drive antibiotic survival as it is the phenotype most transited to and least transited from.

|  | Most frequent transition | Phenotype most transited to | Phenotype least transited from (stable) |
| --- | --- | --- | --- |
| Ciprofloxacin | LS to HS (38%) | HS (38%) | HR (0%) |
| Meropenem | HS to HR (33%) | HR (33%) | LR (3%) |
| Tobramycin | LS to HS and LS to LR (20% each) | HS (35%) | HS (15%) |

**Fig. 4.**
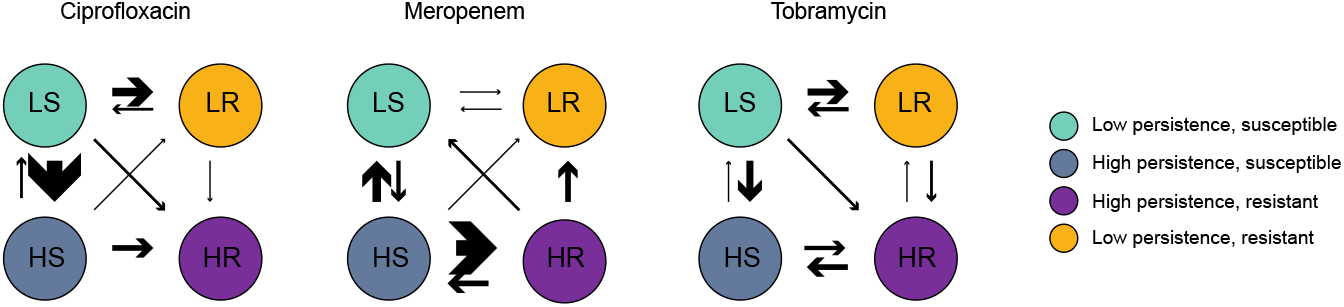
Transitions between the four different responses to antibiotics, identified from phenotypes mapped to the phylogenies. Number of independent events written by corresponding arrows. Panels show all events for DK1 and DK2 together. See Fig. S9 for DK1 and DK2 separately). LS = low persistence | susceptible; LR: = low persistence | resistant; HS: high persistence | susceptible; HR: high persistence | resistant. Arrow thickness denotes number of transitions between categories.

**Fig. 5.**
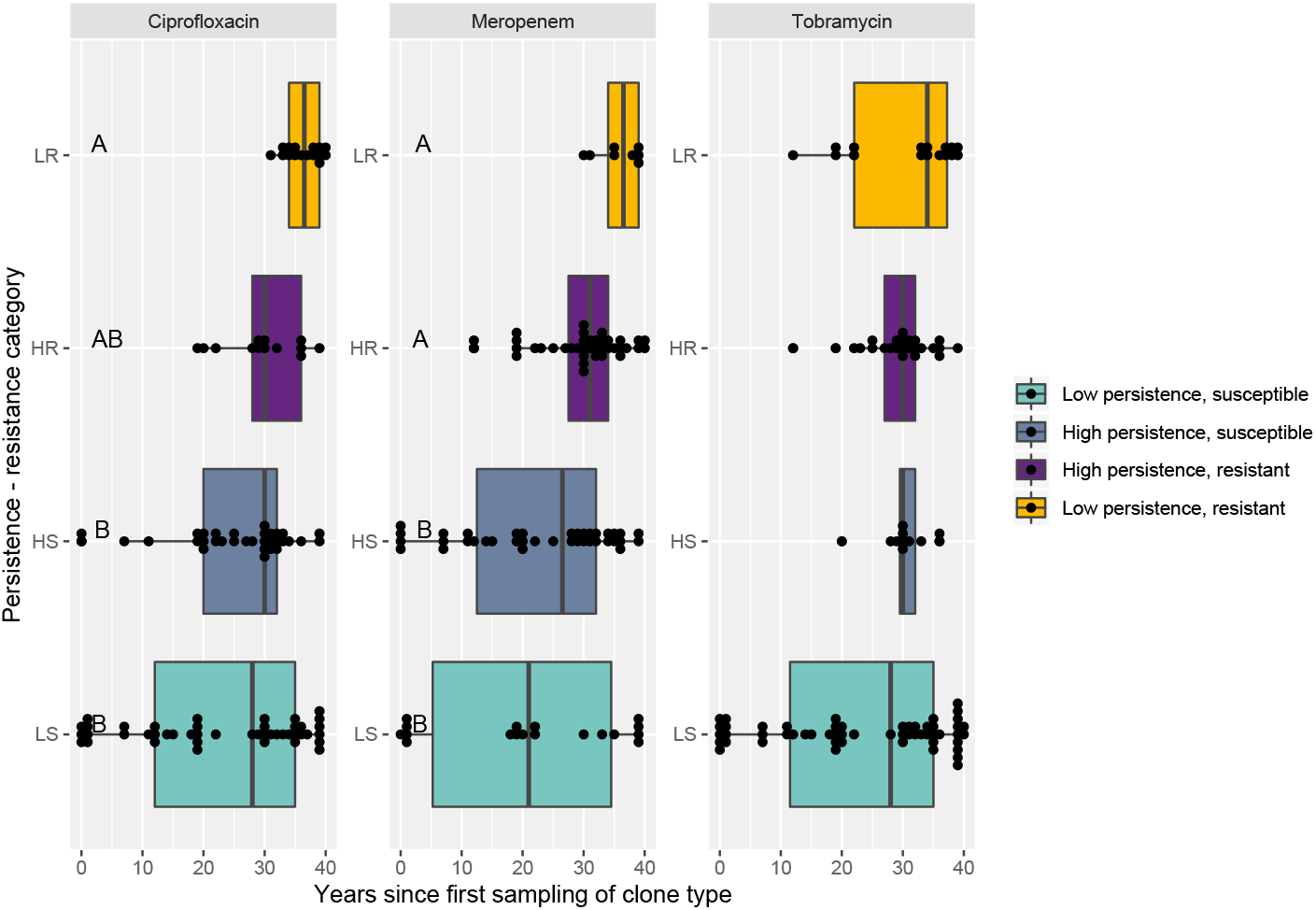
Isolate sampling time for persistence-resistance categories. Boxplots show median sampling time ± 25 percentiles and values as dots. Grouping by 2 -way ANOVA Tukey HSD test p < 0.05 denoted by letters A – C (Table S8). LS = low persistence, susceptible; LR: = low persistence, resistant; HS: high persistence, susceptible; HR: high persistence, resistant. For plots of DK1 and DK2 separately see Fig. S8.

While persistence may readily evolve as a first response, this is frequently followed by resistance, whereas resistance does evolve in the absence of persistence. Resistance and persistence to tobramycin and ciprofloxacin are equally likely to evolve first (ciprofloxacin, LS to LR [P(X ≥ 6) ∼ pois(X; 8.5) > 0.05]; tobramycin LS to LR [P(X ≥ 6) ∼ pois(X; 5) > 0.05], (Fig. 4 & fig. S3A-H; Table S4), while persistence to meropenem already is the ancestral state for DK1. Achieving both resistance and high persistence (HR) is more likely to occur by first developing high persistence, then resistance, compared to resistance first followed by high persistence. This is significant when calculated for all antibiotics together (HS to HR [P(X ≥ 18) ∼ pois(X; 10.5) = 0.02]), while sample sizes are too small to compare for each antibiotic individually.

#### Isolates with low population density have higher levels of resistance and persistence

We find that maximum bacterial density after 24 hours of growth is negatively correlated with both persistence and resistance. This indicates that resistant and persister isolates either grow slower or have an extended lag and/or stationary phase. Resistant isolates that are either high or low persisters consistently have lower population densities (purple and yellow boxes in Fig. 6; Table S7). While the high persister phenotype does occur in isolates of higher density (grey boxes), the high persistent | resistant (HR) phenotype significantly has a lower density in the presence of all three antibiotics (purple boxes). This pattern was in particular driven by the DK1 clonal lineage (Fig. S7). There was, however, also a significant decrease in maximum density with infection time despite the resistant or persistent phenotypes (Linear regression; DK1: R^2^ _adj_ = 0.50, p < 0.001; DK2: R^2^ _adj_ = 0.21, p < 0.001). So the negative correlation between resistance and high persistence with low density could be confounded by the fact that resistant isolates tend to be sampled late in infection. Analysing the data including both sampling time and population density to test for their effect on the evolution of resistance and persistence, shows that lower population density in itself is associated with an evolved antibiotic response (except for persistence to meropenem treatment in DK2; Table S5 & S6).

**Fig. 6.**
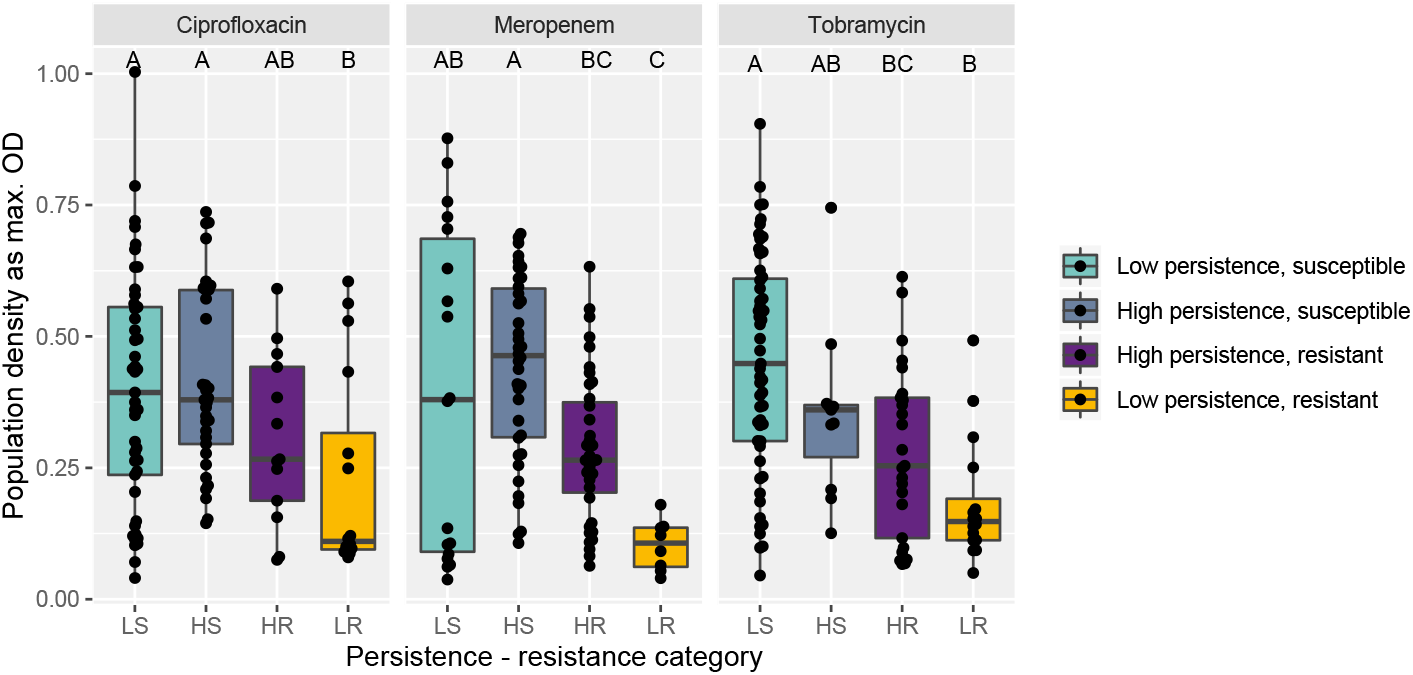
Population density measured as max OD for persistence-resistance categories. Boxplots show median max OD ± 25 percentiles and values as dots. Grouping by 2 -way ANOVA Tukey HSD test p < 0.05 denoted by letters A – C, groups are significantly different if they do not share a letter (Table S7). LS = low persistence | susceptible; LR: = low persistence | resistant; HS: high persistence | susceptible; HR: high persistence | resistant. For plots of DK1 and DK2 separately see Fig. S7.

## Discussion

We investigated the evolution of antibiotic persistence and resistance during pathogen evolution. We find that both persistence and resistance to three clinically relevant antibiotics, meropenem, ciprofloxacin and tobramycin, are selected for during infection. However, we observe differences in antibiotic response strategies between clone types and type of antibiotic used. While persistence and resistance are equally likely to evolve first, persistence can act as a stepping stone to gain resistance. In contrast, we did not find that resistance acted as a stepping stone for persistence. As such, our *in situ* results are in line with recent findings by experimental evolution of *P. aeruginosa* (10) and *E. coli* (21, 22). In our collection of isolates, we find both strategies take time to evolve; persistence evolves *de novo* after 7-19 years and clinical resistance after 12-20 years. This suggests that other mechanisms such as biofilm production or limited diffusion of antibiotics within the CF lung must play a role for the establishment of pathogens despite antibiotic treatment early in infection (13, 40, 41).

We find that individuals are typically colonized by antibiotic susceptible and non-persistent phenotypes, and that strategies to evade the effect of antibiotics are, therefore, unsurprisingly favored by selection in the lung (Fig. 1 -3). The pattern of how these traits spread over time differs, however, between the two transmissible lineages we examined. One of the lineages, DK2, shows a significant increase in persistence over time in response to all three antibiotics, while DK1 only shows increased persistence under tobramycin treatments, and significant decrease in persistence under meropenem and ciprofloxacin treatments. The differences between DK1 and DK2 may reflect topology of the phylogenies and ancestral state: the DK1 phylogeny has several distinct subclades that evolve independently (Fig. S1A). Some clades maintain “wild-type-like” production of virulence factors after more than 30 years of infection (e.g. the siderophore pyoverdine (42) and protease (30)). The ancestral state is high persistence for DK1 in the meropenem treatment, and high persistence under ciprofloxacin evolves nine times independently (Fig. S3A & S9), first in 1973, the first year of sampling (Fig. 1). After 40 years of infection, some isolates still exhibit low persistence or susceptibility to one antibiotic, however, very few are susceptible and low persisters to *all* antibiotics (DK1: seven isolates sampled from 1992-2012; DK2: one isolate sampled in 1992). This perhaps reflects variation in selection pressure due to differences in drug treatment between individuals and/or infections in different areas of the CF lung (30, 43).

An outstanding question is how specific persistence is to the type of antibiotic that cells are exposed to. We found a significant overlap in persistence phenotypes between ciprofloxacin and meropenem but not with tobramycin. In contrast, a recent study (10) shows correlations of the persister phenotype between ciprofloxacin and tobramycin. This may be because the authors select only a subset of strains from a lineage of susceptible strains with MIC values below the clinical breakpoint and use much lower doses of antibiotics relative to the MIC than in our assays (approximately 10-100 fold lower). Our results may reflect that ciprofloxacin and meropenem were simultaneously used in the clinic for treatment, exerting comparable selection pressures. Another explanation could be that despite selection for the canonical antibiotic resistance genes, exposure to one antibiotic could select for genome wide mutations in metabolic genes that can contribute to higher than MIC resistance to multiple antibiotics (5, 44). Additionally, the CF lung is very complex and many other factors may be selecting for persistence, such as evasion to the immune system and exposure to a stressful oxidative environment (17).

We also found similarity between the response to ciprofloxacin and meropenem is that some isolates from early infection with a high persistence phenotype exhibit a “revival” phenotype to meropenem and/or ciprofloxacin. This is defined as no growth observed in the undiluted cultures, however, once the antibiotics are diluted 10-fold (Fig. S5), growth is observed (isolates shown as light red circles in Fig. S1). This phenotype indicates the presence of persisters that are only detected when the antibiotic is removed from their environment. This “revival” phenotype is only observed in the early isolates, and we speculate that this could be a potential intermediary route to evolve high persistence. All other isolates classified as high persister had high CFU counts from both undiluted and 10-fold diluted cultures. The correlation between meropenem and ciprofloxacin could be shaped by the antibiotic killing mode and bacterial resistance mechanisms: *P. aeruginosa* cells can tolerate ciprofloxacin exposure by elongating without dividing, allowing cells to survive longer in the antibiotic such that when it is removed they can divide and grow (45-47). Meropenem can be enzymatically degraded by beta-lactamase activity (48). This detoxifies the environment from meropenem at a faster pace than an antibiotic which is not, allowing cells to revive faster (49). In contrast, tobramycin tends to be bactericidal to *P. aeruginosa*, it disrupts bacterial cells membranes killing cells upon exposure to high concentrations (50), such that cells either persist or die with no revival intermediary. This may also explain why we observe fewer high persister isolates in the presence of tobramycin treatment (Fig. 6).

We find that maximum bacterial density is negatively correlated with both persistence and resistance. This indicates that persisters either grow slower or have an extended lag and/or stationary phase; both traits that have been observed to contribute to persistence (51-55), and a common phenotype in chronic infections (40). This is in accord with the notion that persisters are dormant cells with reduced metabolism and/or non-dividing that are also found in deep biofilm layers (13). It has also been shown that resistance often comes at a metabolic cost that may lower growth rate (56, 57). Additionally, because later stages of infection, in our study, are characterised by isolates that achieve smaller populations densities in vitro but higher persister counts, this indicates that the fraction of persisters in the population will be proportionally higher in these compared to isolates with a higher maximum density. However, it is important to note that slow growth is a common phenomenon in *P. aeruginosa* isolates from late stage infection and this may be influenced by factors other than antibiotic response strategies.

Persistence is a complex polygenic trait, that may on one hand be under selection by variables other than antibiotics, such as oxidative stress and host immune evasion, and on the other hand also be affected by selection on other phenotypes such as growth rate. Therefore, we here focus on the phenotype, and not the genotype, of persistence. While most work has focused on resistance, we are with this work achieving a better understanding of the correlation of persistence to resistance; how this is coupled to bacterial growth rate; and associations with specific antibiotics. This is important because the discovery of new antibiotics has stalled and our best approach is to use the already available drugs in novel ways – by reconsidering doses and combinations (58, 59). To do so we must continue to explore how bacteria respond and adapt to drugs *in situ*.

In conclusion, our results indicate that persistence and resistance phenotypes rarely evolve before a decade post infection. As expected, our results are consistent with resistance being the primary strategy for surviving antibiotic treatment in late stage chronic infections. Persistence, however, occurs earlier in infection than resistance, when it could be used as a stepping stone to resistance and thereby contribute to the development of recalcitrant infections. Our study highlights the importance of not generalising when drawing conclusions from studies on single clones and/or single antibiotics to better understand persistence and preventing its evolution as a means to mitigate resistance. We cannot assume every infection follows the same course, but that each is influenced by the pathogenic clone type, antibiotic treatment and the host environment. Critical to achieve this is the development of methods for easy detection of persistence, which is not routinely screened for. These results could have implications for early intervention treatment strategies to prevent evolution of persistence and ultimately resistance, e.g. using lower doses of antibiotics and in specific combinations, allowing us to prolong the use of antibiotics for treating infections (58, 59).

## Supplementary information

### See Excel datasheet - for all tables

Table S1: An overview of the non-lung, acute and chronic infecting isolates, and CFU and MIC data. Data given as CFU undiluted (CFU1), diluted 10 fold (CFU10) and diluted 100 fold (CFU100) and the standard error for the replicates for each. Persistence phenotype listed as high/low and high/low/revival. Time from first sampling of clone type to sampling of the specific isolate given as time first.

Table S2: An overview of isolates of the transmissible clone types DK1 and DK2 and CFU and MIC data. Data given as CFU undiluted (CFu1), diluted 10 fold (CFU10) and diluted 100 fold (CFU100) and the standard error for the replicates for each. Persistence phenotype listed as high/low and high/low/revival. Persistence-resistance phenotype listed as LS, HS, HR. time from first sampling of clone type to sampling of the specific isolate give as time first.

Table S3: List of transitions for DK1 and DK2 persistence comparing antibiotics pairwise. Transitions are mapped on Fig. S3G and H, and the table lists to which antibiotic persistence changes, and whether this leads to a match in persistence to the other antibiotic.

Table S4: List of transitions for DK1 and DK2 persistence and resistance phenotypes. Clone_lineage refers to the transition marked on phylogenies in figs. S3A-F. For DK2 branch name is noted.

Table S5: Results from GLMs for antibiotic persistence, testing the effect of time and max OD, and the interaction between the two. In blue is highlighted p-values > 0.05 and the best fit model with the lowest Akaike information criterion (AIC). Analysis for DK1 and DK2 together, and separately.

Table S6: Results from GLMs for antibiotic resistance, testing the effect of time and max OD, and the interaction between the two. In blue is highlighted p-values > 0.05 and the best fit model with the lowest Akaike information criterion (AIC). Analysis for DK1 and DK2 together, and separately.

Table S7: Results from 2-way ANOVA, testing the effect of max OD on persistence and resistance, and the interaction between the two. Analysis for DK1 and DK2 together, and separately.

Table S8: Results from 2-way ANOVA, testing the effect of length of infection on persistence and resistance, and the interaction between the two. Analysis for DK1 and DK2 together, and separately.

**Fig. S1.**
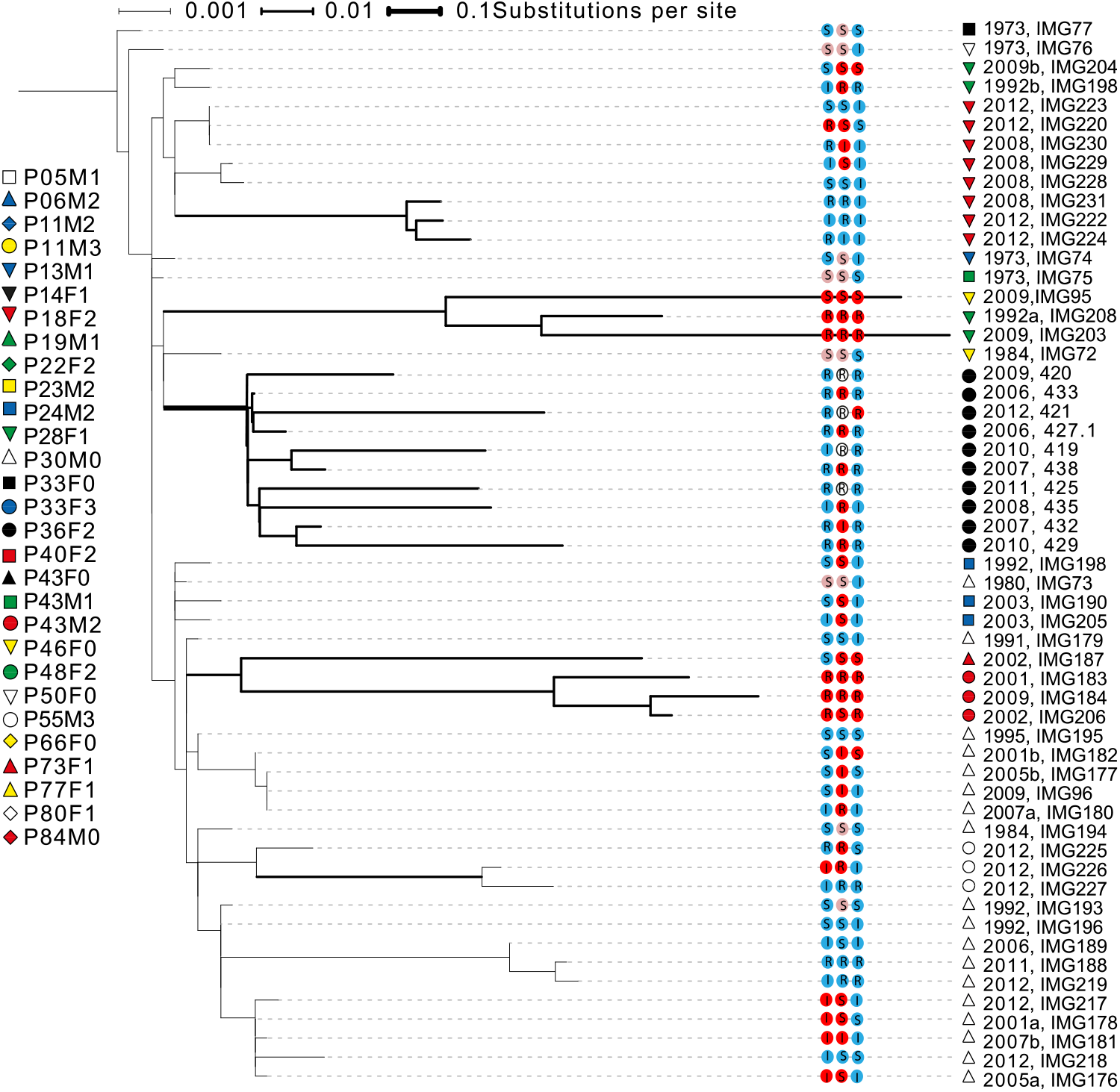
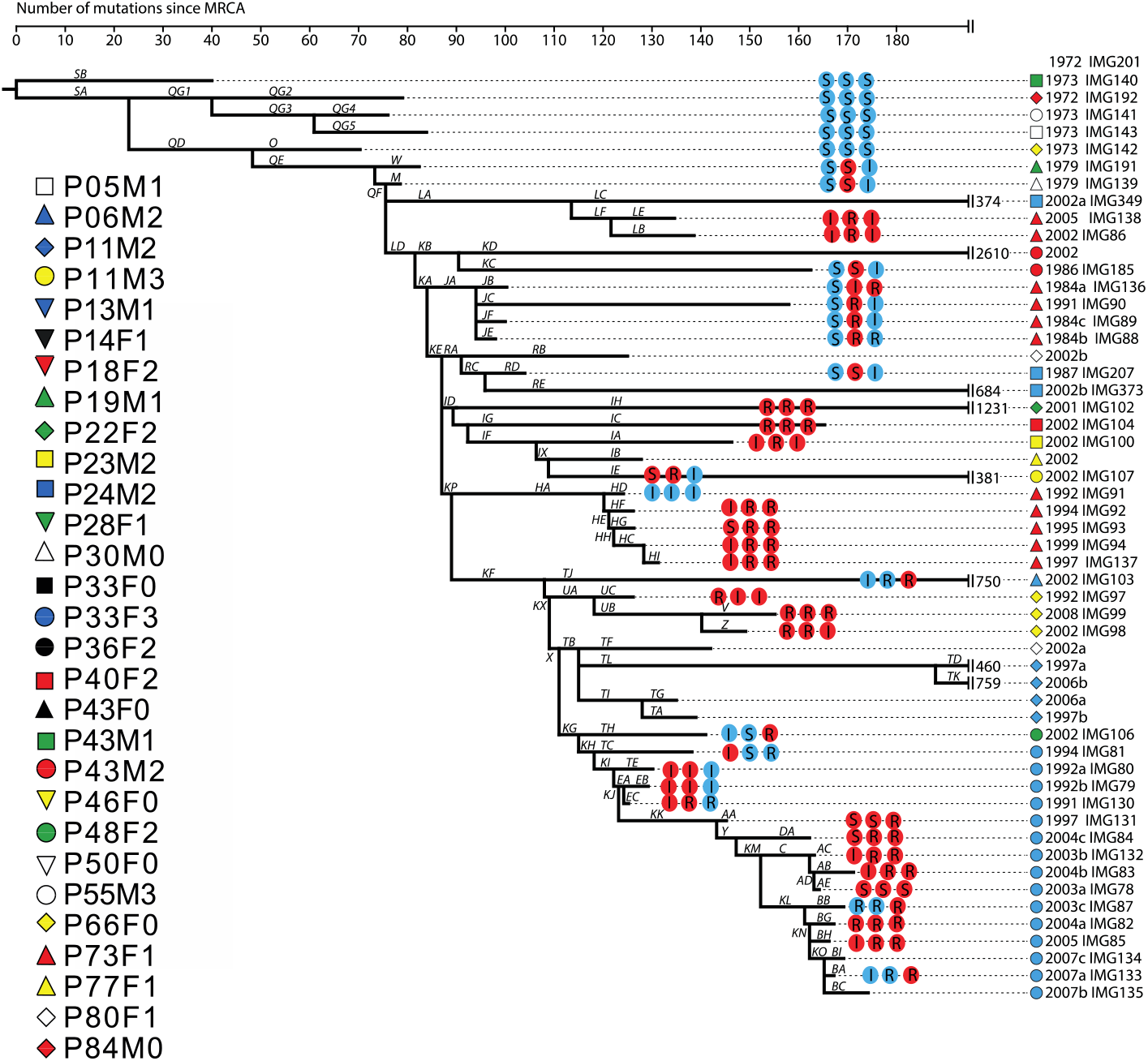
A & B Phylogenies of DK1 and DK2, modified from Andersen et al. 2019 (DK1) and Marvig et al. 2013 (DK2). For each isolate, persistence under treatment with the three antibiotics (in the order ciprofloxacin, meropenem, and tobramycin) is shown in blue (low) or red (high). and marked as susceptible (S), intermediate (I) or resistant (R) to the antibiotics. Isolates that only show high persistence when the culture is diluted 10 times are shown in pink (only found in DK1).

**Fig. S2.**
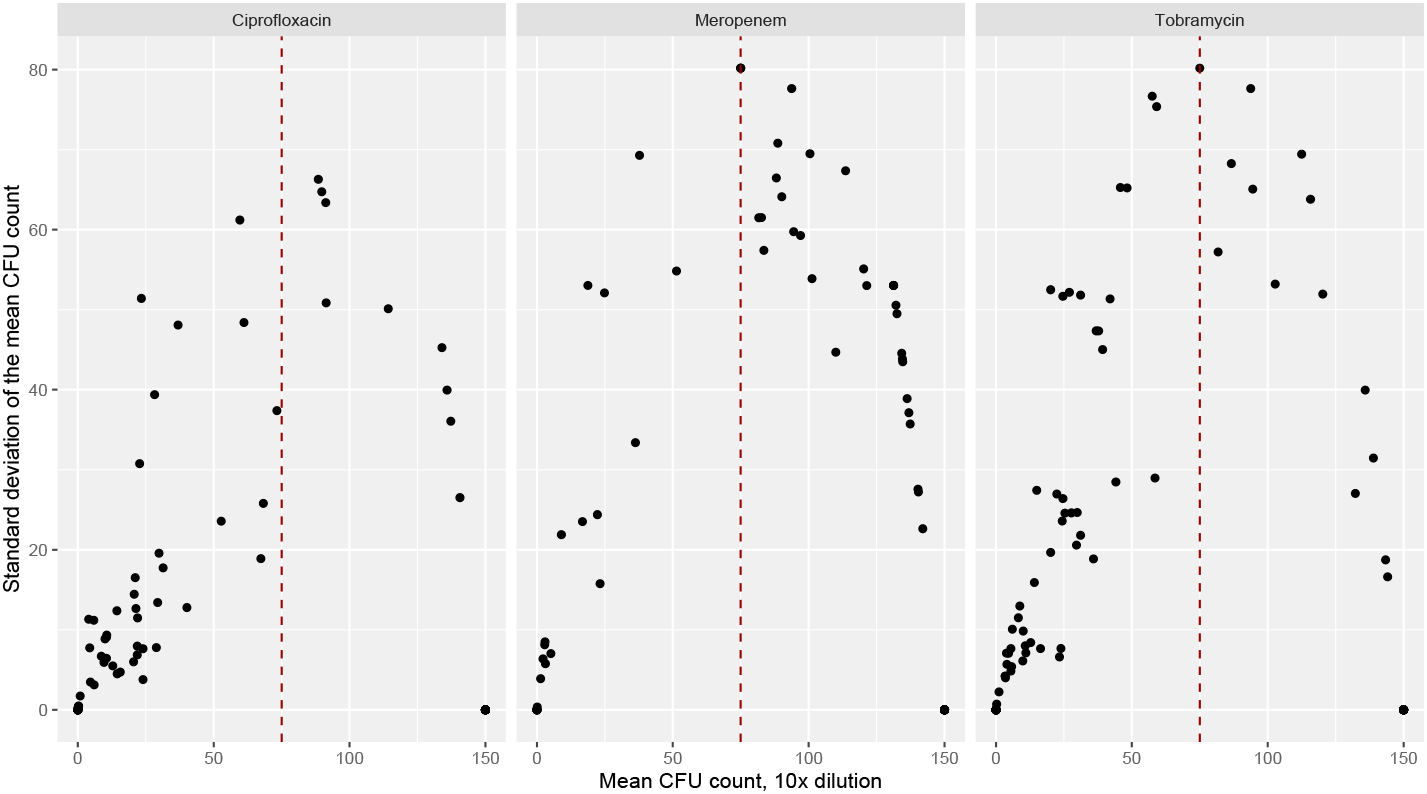
Correlation between the mean CFU count at 10 times dilution, and the standard deviation around the mean. There is a bell-shaped association, as isolates with an intermediate count tend to represent isolates with replicates of either very high or very low counts. The dashed red line marks the cut-off of 75 CFU to classify isolates as either low or high persisters.

Fig. S3: Phylogenies of DK1 and DK2 with the transitions in persistence and antibiotic resistance to three different antibiotics mapped. A: DK1 persistence and resistance, ciprofloxacin; B: DK1 persistence and resistance, meropenem; C: DK1 persistence and resistance, tobramycin; D: DK2 persistence and resistance, ciprofloxacin, E: DK2 persistence and resistance, meropenem, F: DK2 persistence and resistance, tobramycin; G: DK1 persistence, all three antibiotics; H: DK2 persistence, all three antibiotics (see extra uploaded files).

**Fig.S4.**
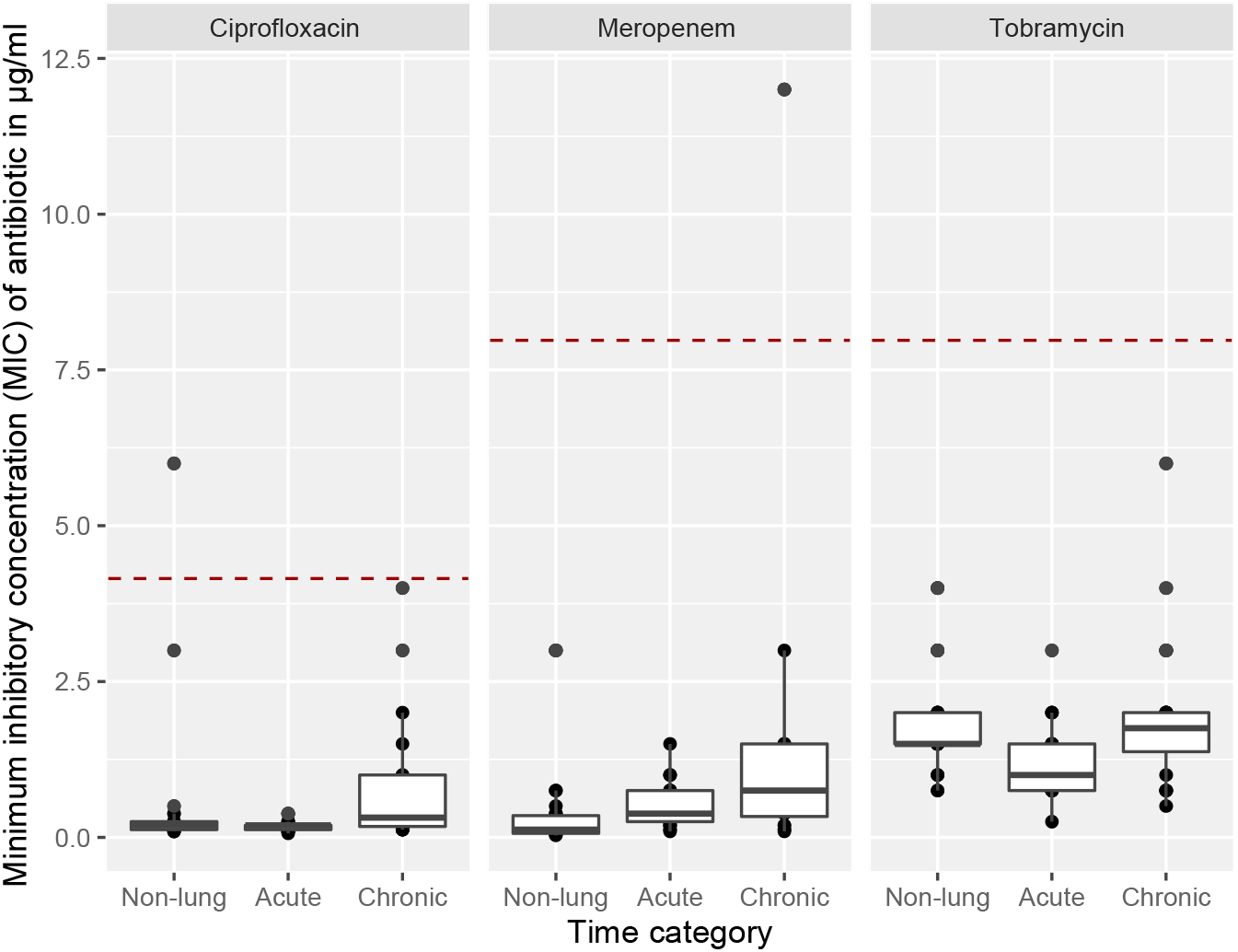
Antibiotic resistance as the minimum inhibitory concentration MIC of antibiotic for each isolate across the non-lung and chronic infecting categories. Boxplots show media MIC ± 25 percentiles and values as dots. Dashed red lines indicate clinical cut-off for classification of resistance (MIC of 4 μg/ml for ciprofloxacin, and 8 μg/ml for meropenem and tobramycin). All except one isolate in the late category for meropenem are in the susceptible category as per clinical cur off limits.

**Fig. S5.**
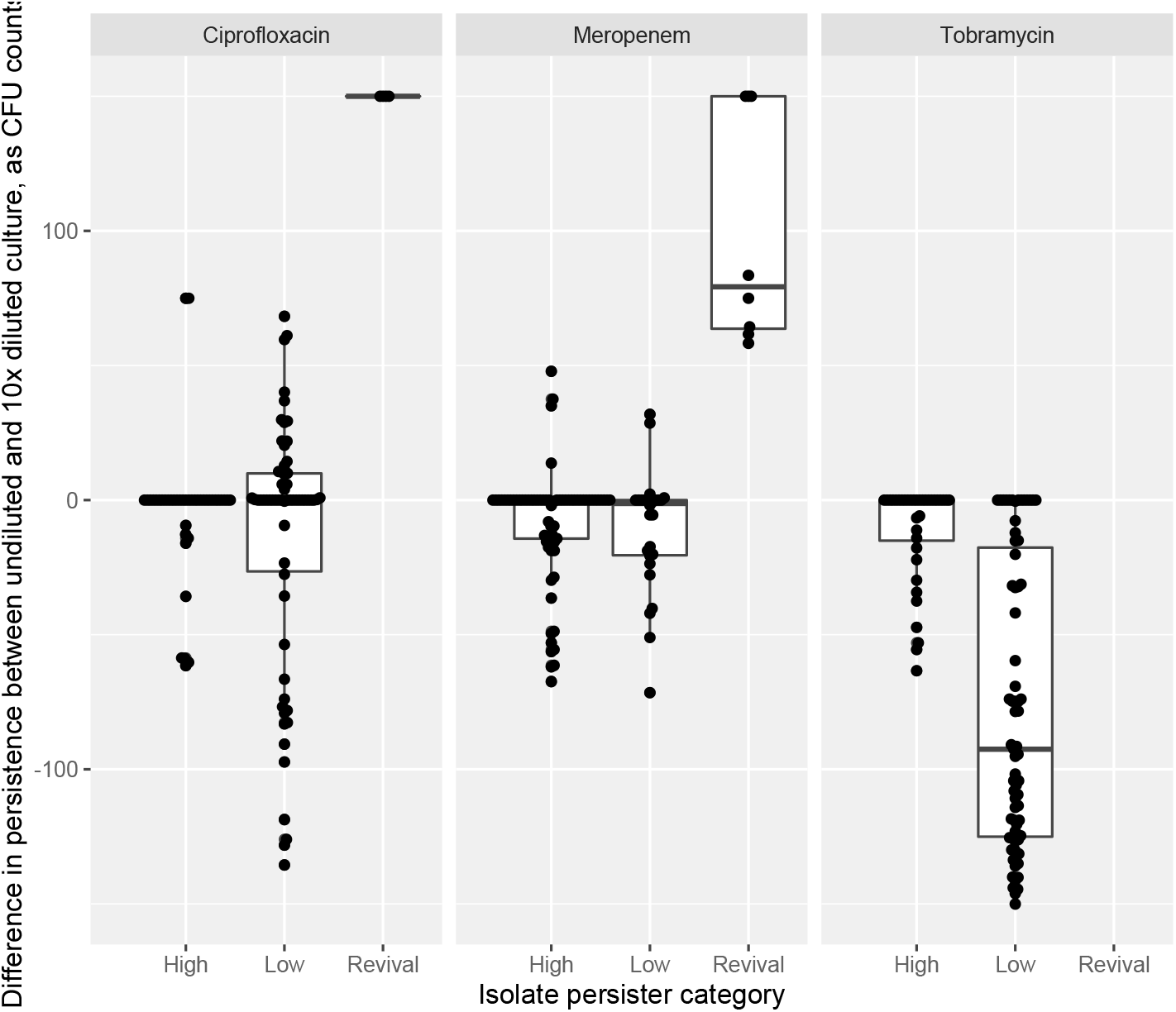
Graphs show the difference in persistence measured as CFU counts from undiluted culture or culture diluted 10 times for isolates classified as low persisters, high persisters, and isolates that “revive” and become high persisters only when antibiotic is diluted. Only the “revival” category has a mean difference above 50. Boxplots show median CFU counts ± 25 percentiles and values as dots.

**Fig. S6.**
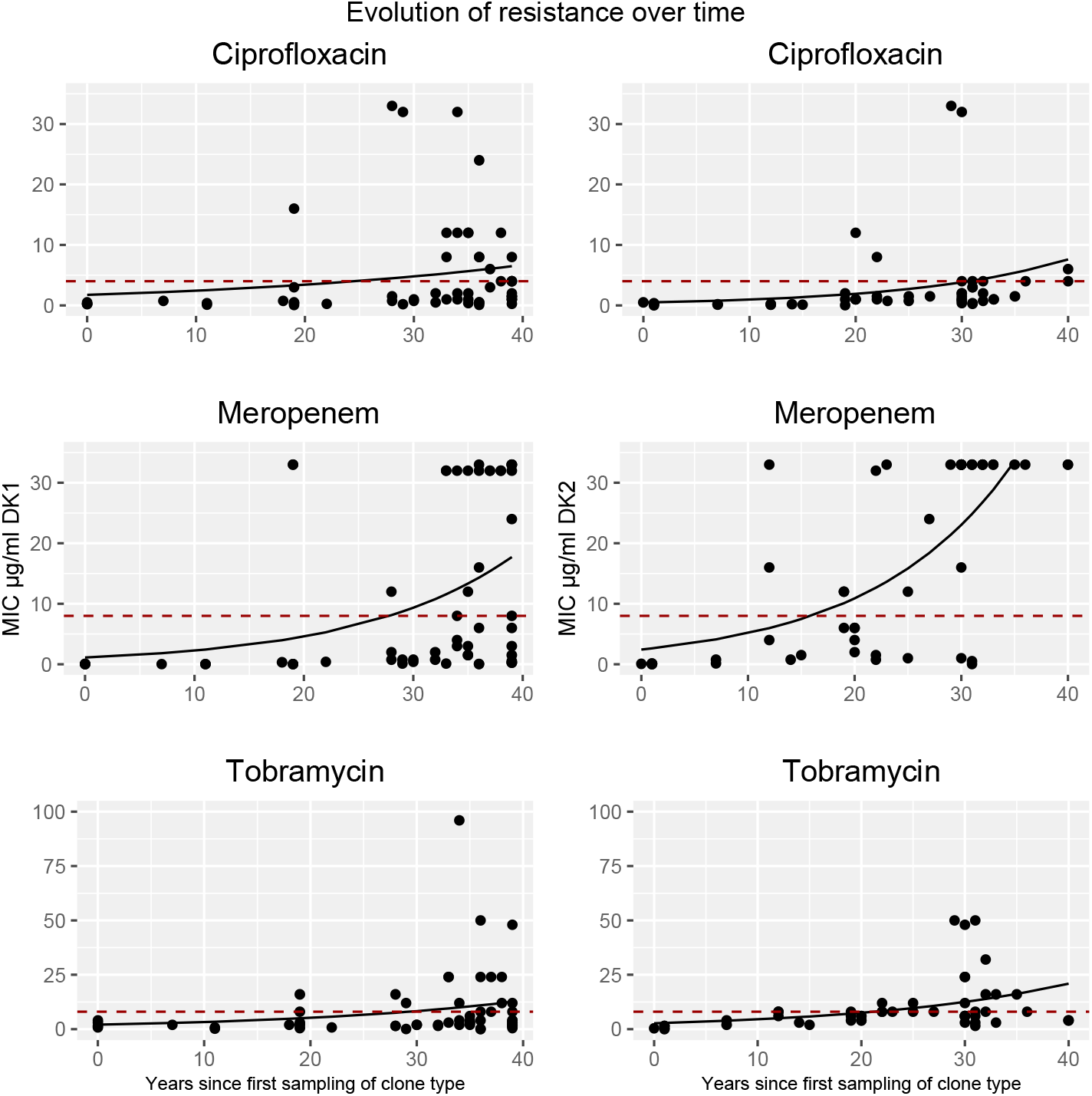
Graphs show resistance, measured as MIC, over time split by clone type, (DK1 left panels, DK2 right panels) for three different antibiotics, with the GLM fits. Dashed red lines indicate cut-off for classification of resistance (MIC of 4 for ciprofloxacin, and 8 for meropenem and tobramycin). Note difference on y-axis scale for the two clone types for meropenem and tobramycin.

**Fig. S7.**
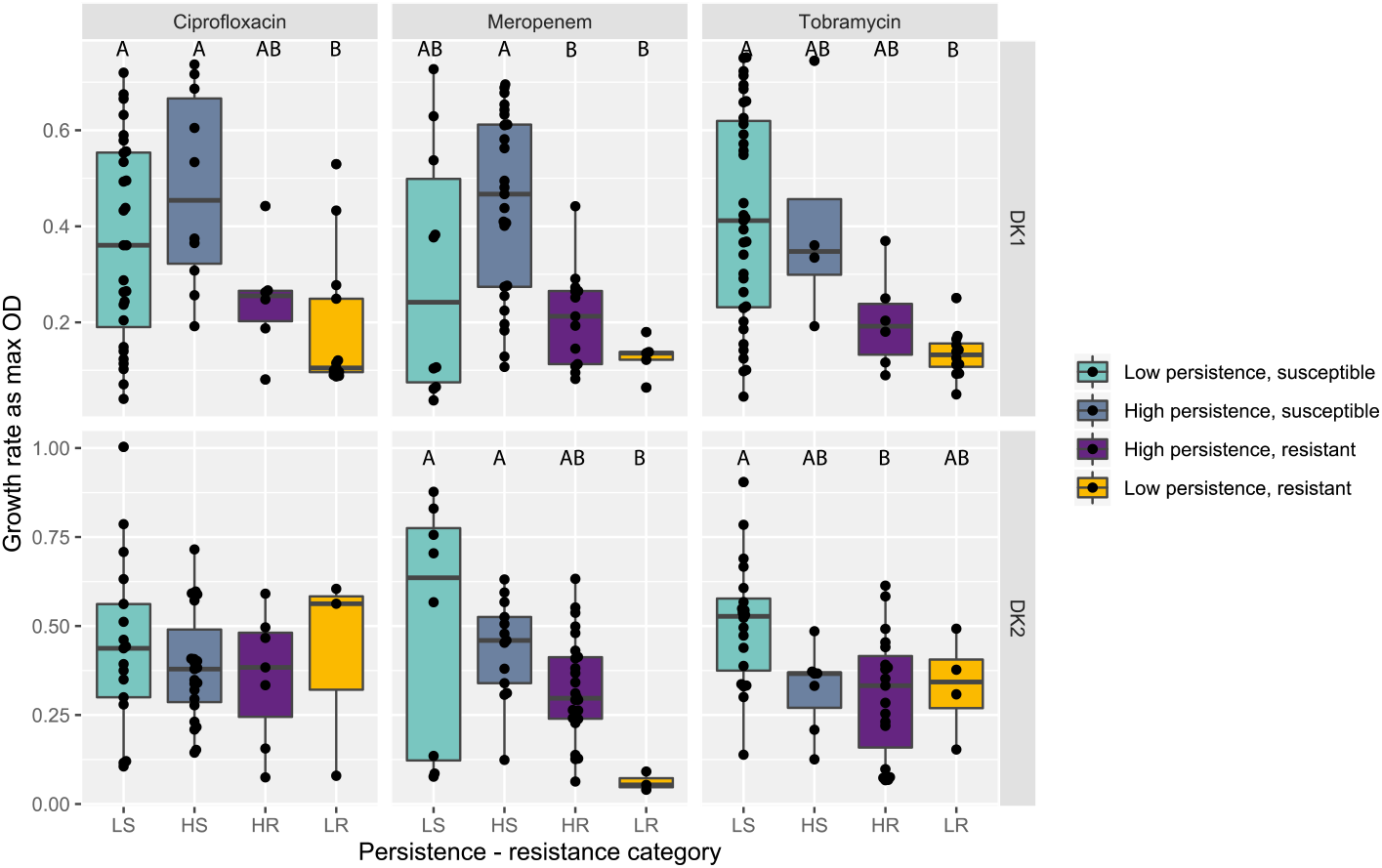
Population density measured as max OD for persistence-resistance categories for the three antibiotics split by clone type, DK1 in top panel, DK2 below. Boxplots show median max OD ± 25 percentiles and values as dots. Grouping by 2-way ANOVA Tukey HSD test p < 0.05 denoted by letters A – C, groups are significantly different if they do not share a letter (Table S7). LS = low persistence, susceptible; LR: = low persistence, resistant; HS: high persistence, susceptible; HR: high persistence, resistant.

**Fig. S8.**
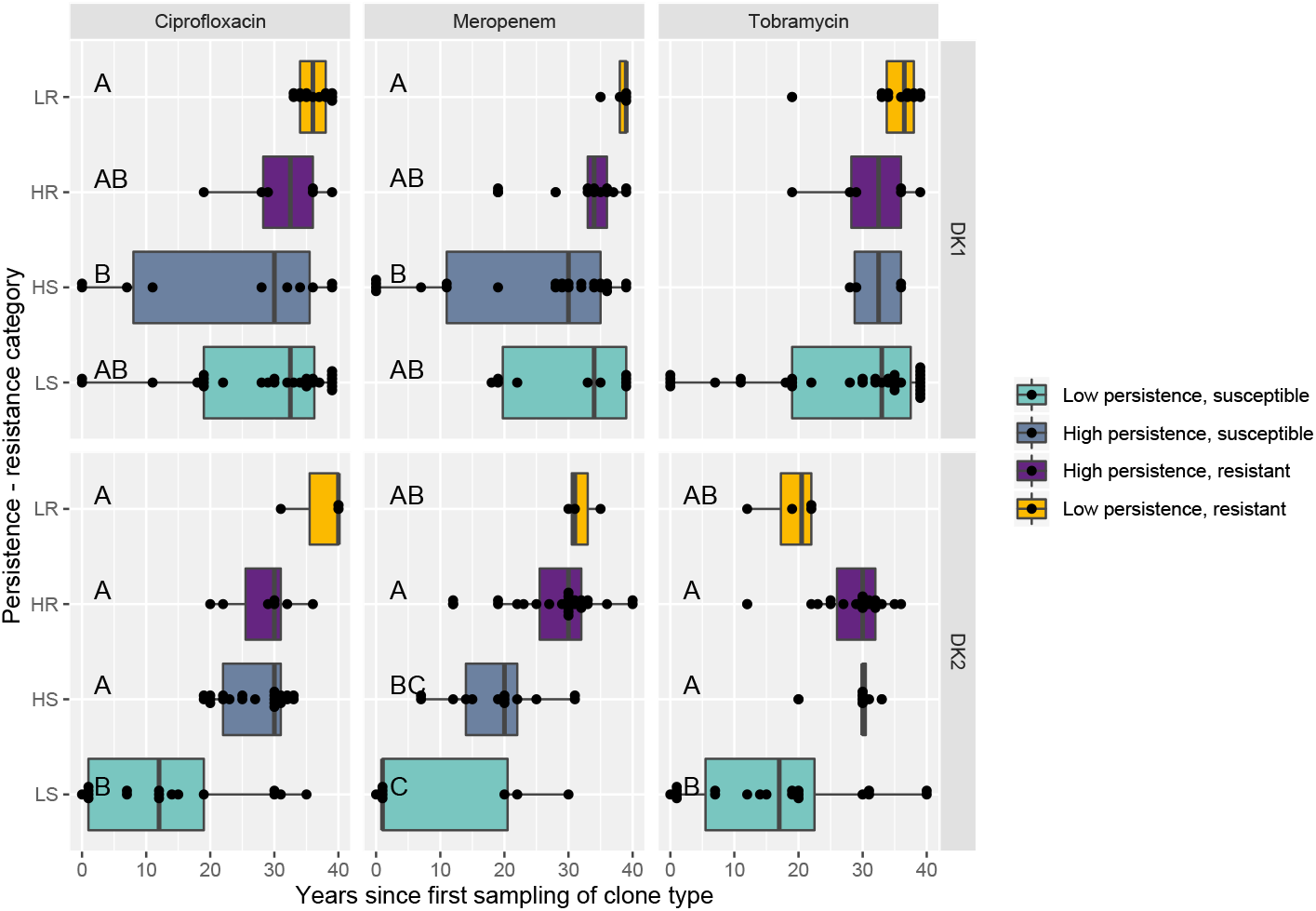
Isolate sampling time for persistence-resistance categories for the three antibiotics split by clone type, DK1 in top panel, DK2 below. Boxplots show median sampling time ± 25 percentiles and values as dots. Grouping by 2-way ANOVA Tukey HSD test p < 0.05 denoted by letters A – C, groups are significantly different if they do not share a letter (Table S8). LS = low persistence, susceptible; LR: = low persistence, resistant; HS: high persistence, susceptible; HR: high persistence, resistant.

**Fig. S9:**
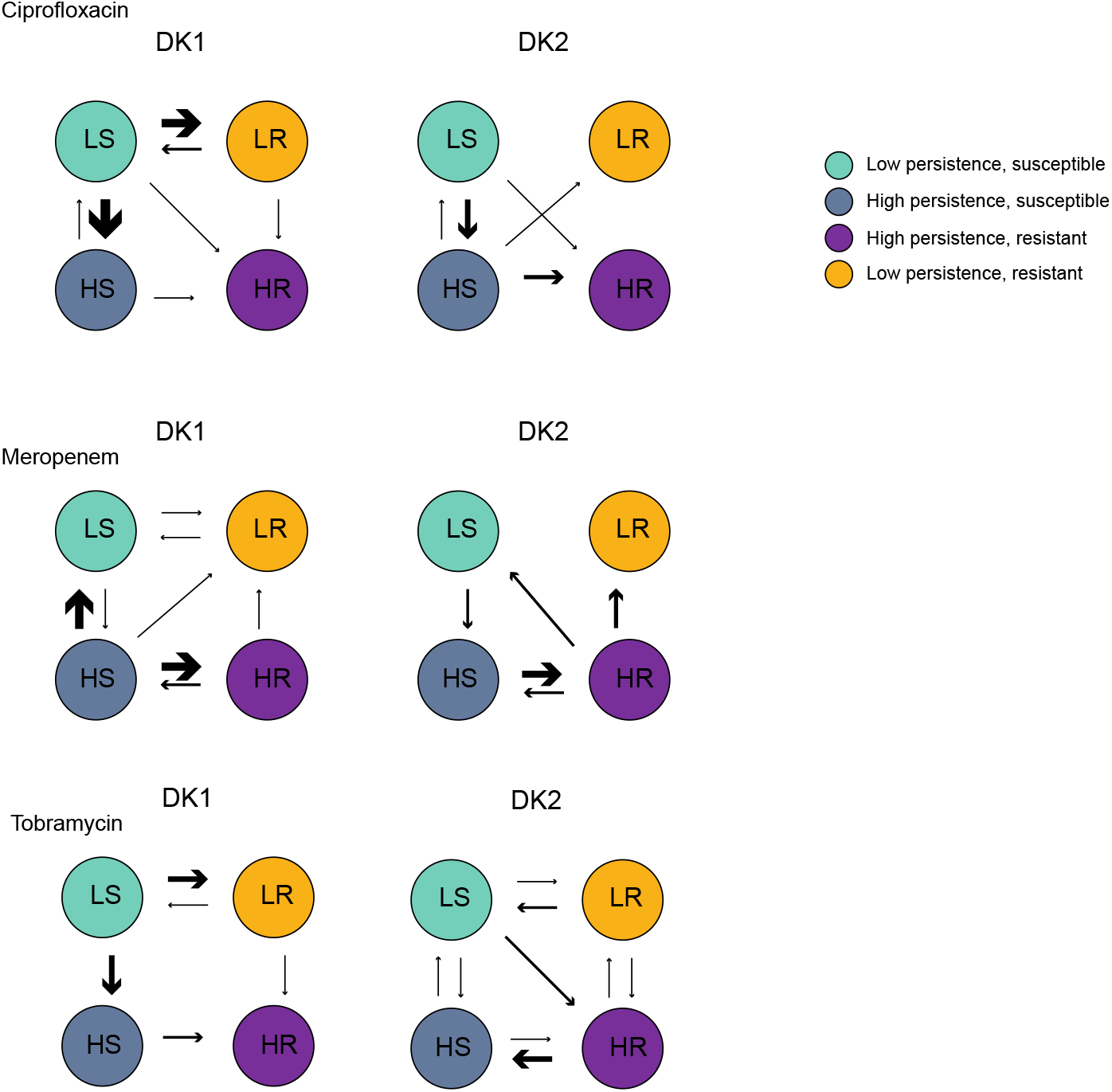
Transitions between the four different responses to antibiotics, identified from phenotypes mapped to the phylogenies of DK1 and DK2. Number of independent events written by corresponding arrows. LS = low persistence, susceptible; LR: = low persistence, resistant; HS: high persistence, susceptible; HR: high persistence, resistant. Increasing arrow thickness denote increasing transitions between categories.

The authors claim no competing interests

---

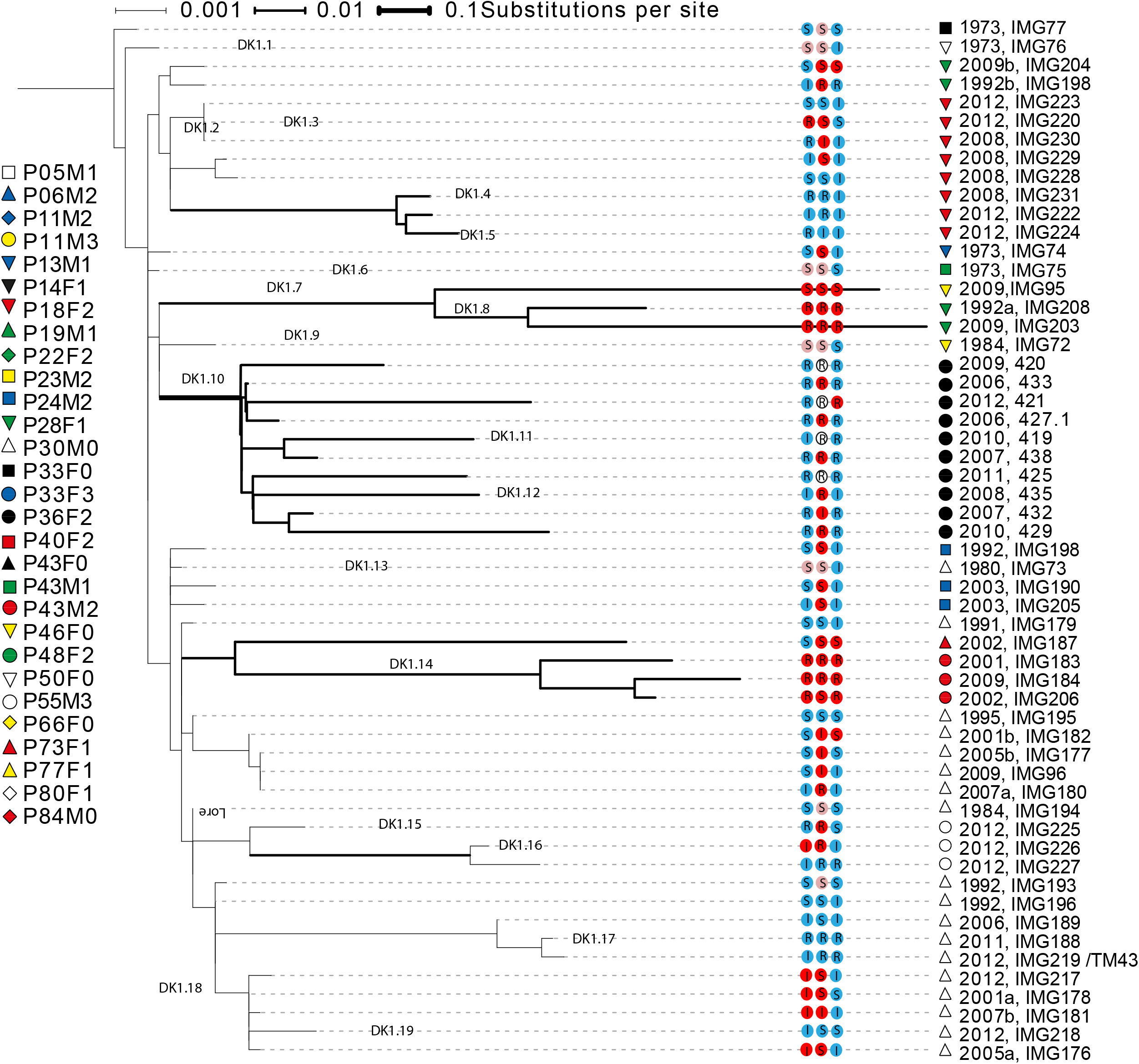

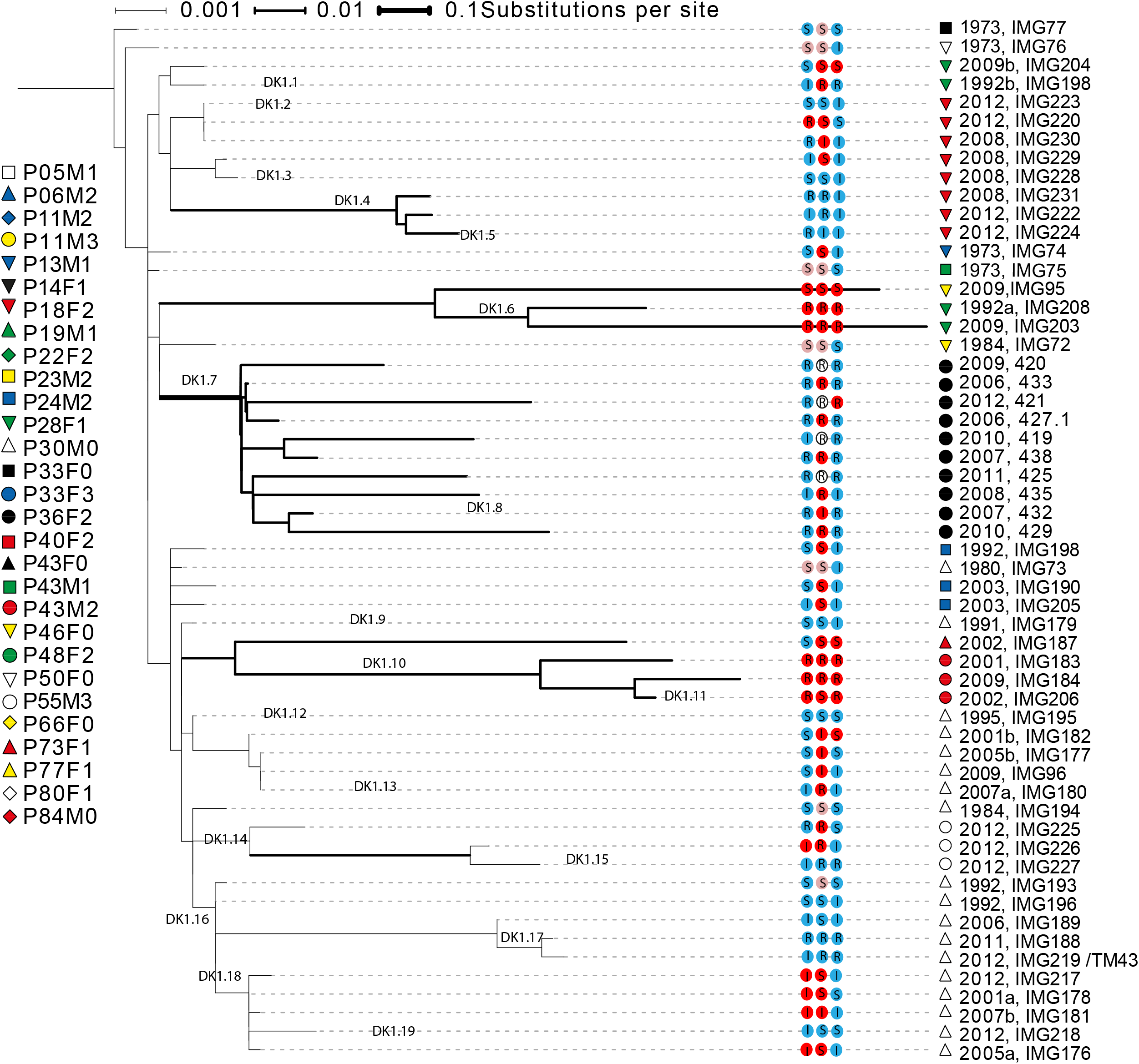

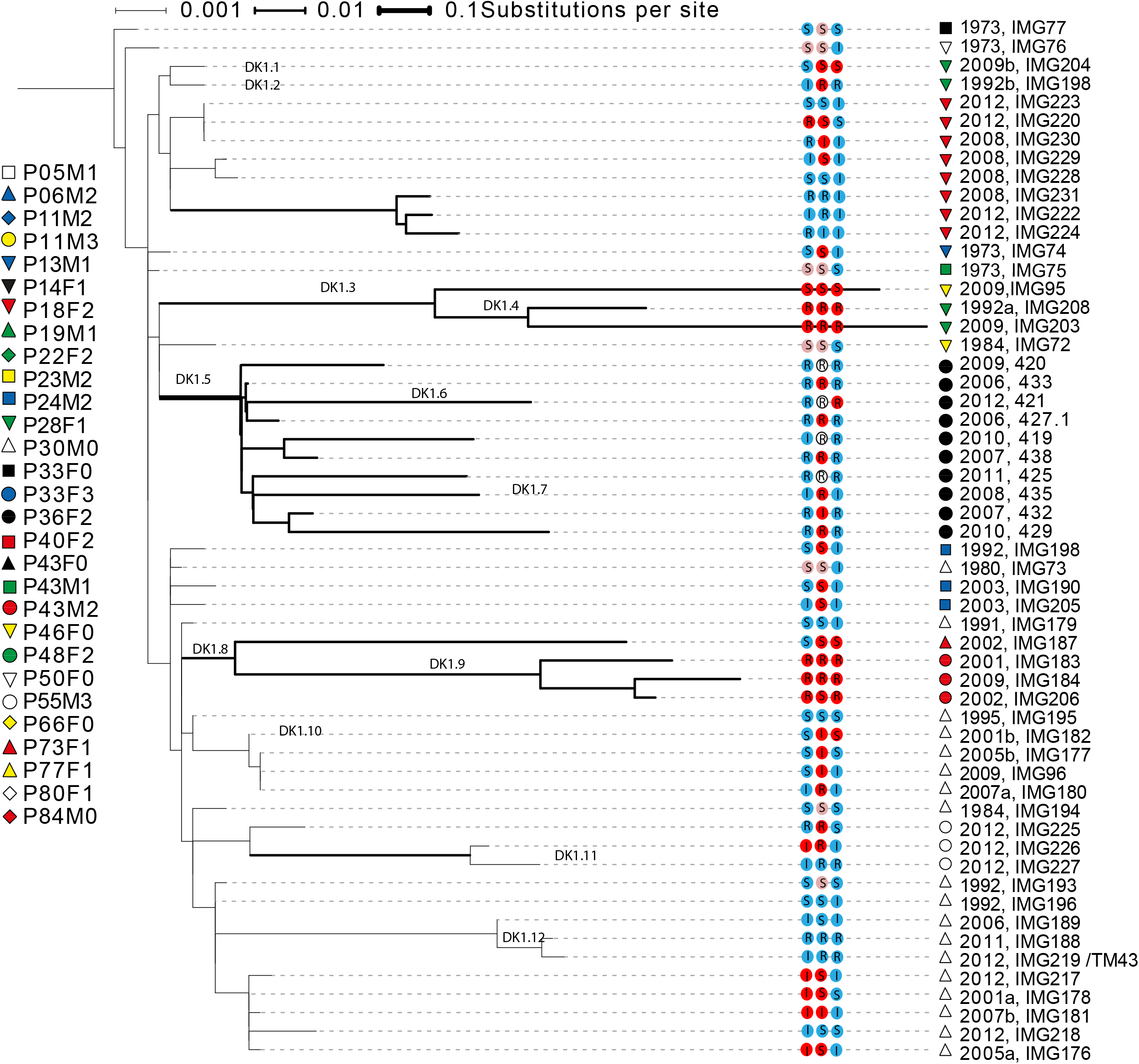

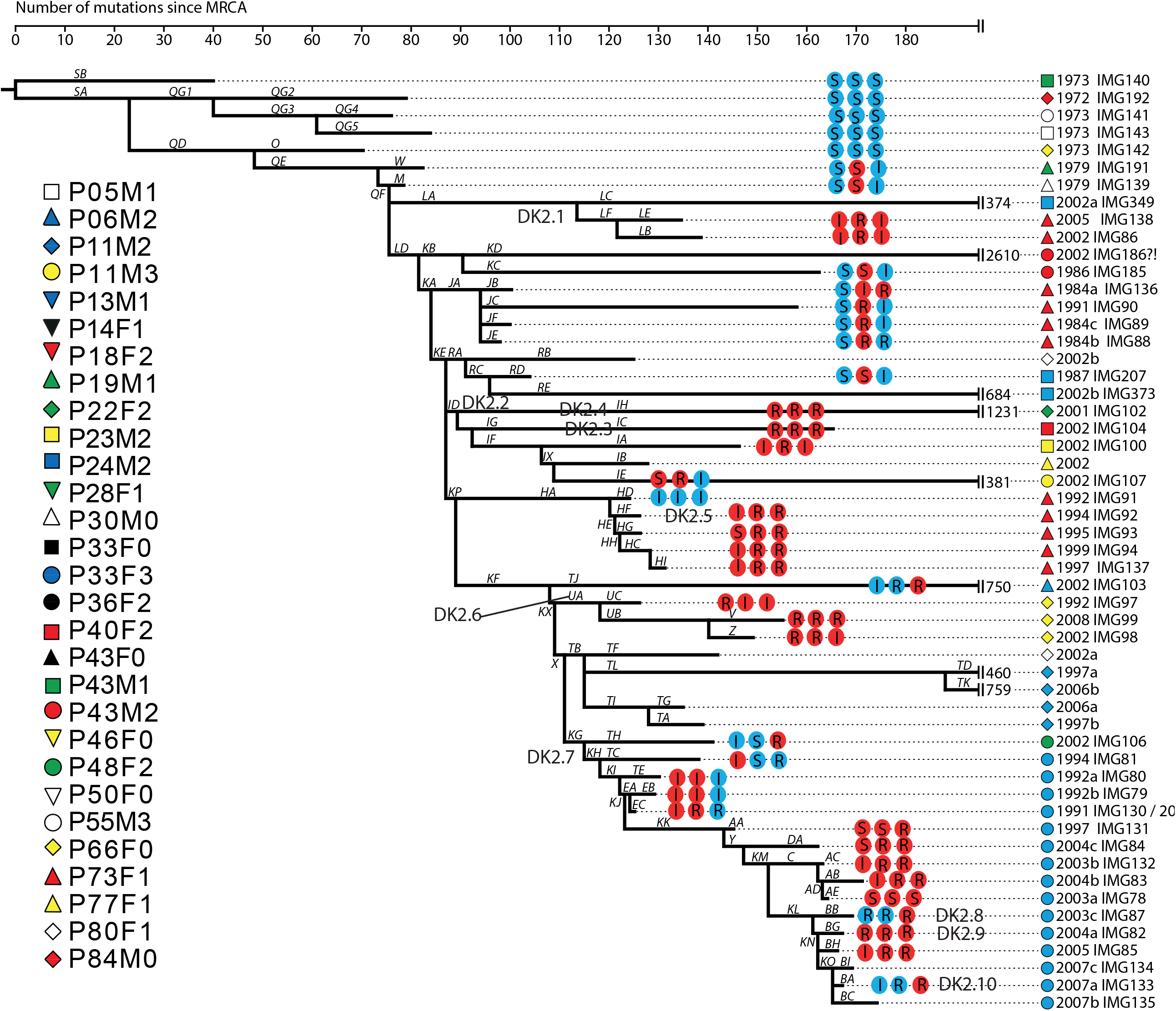

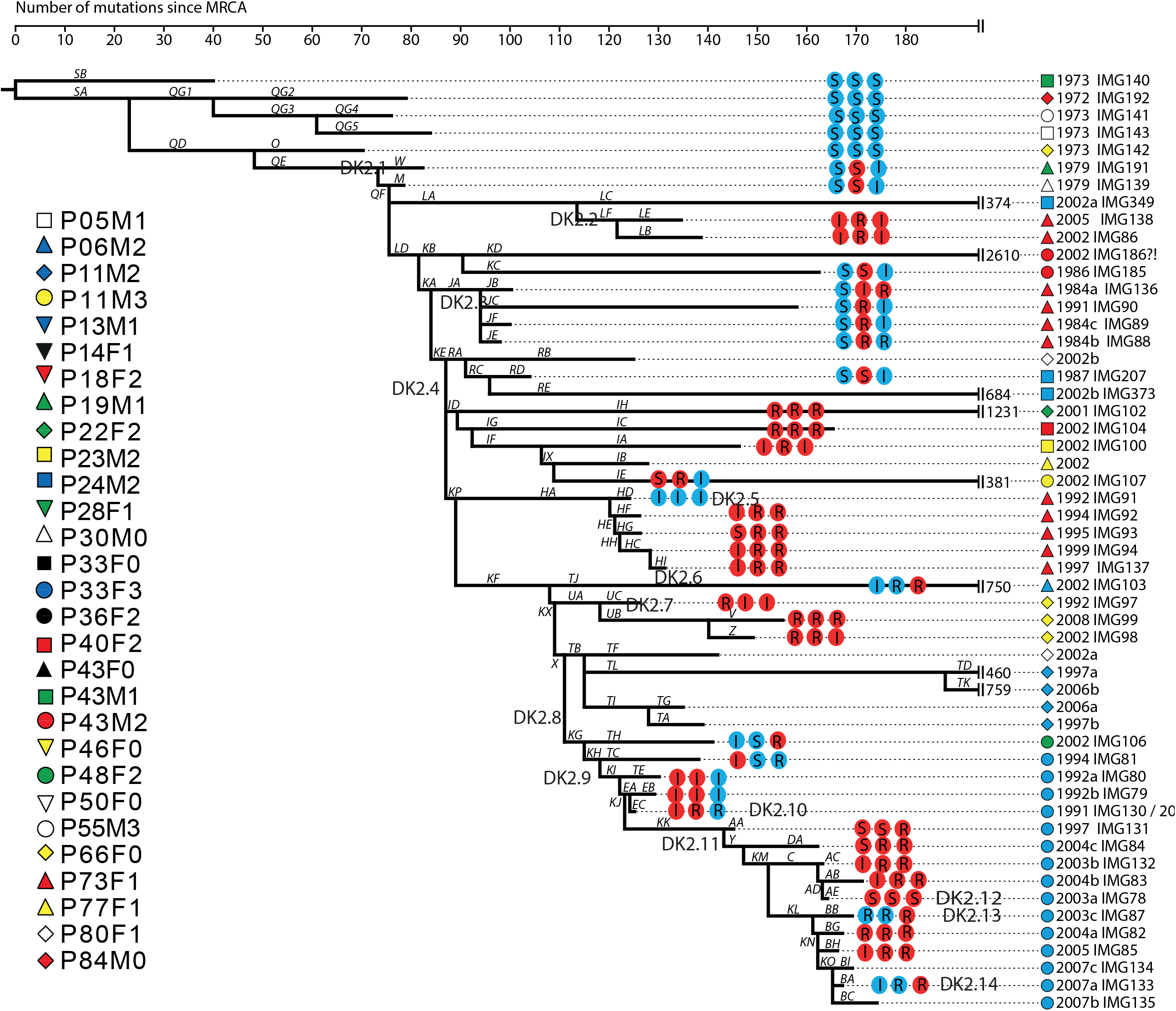

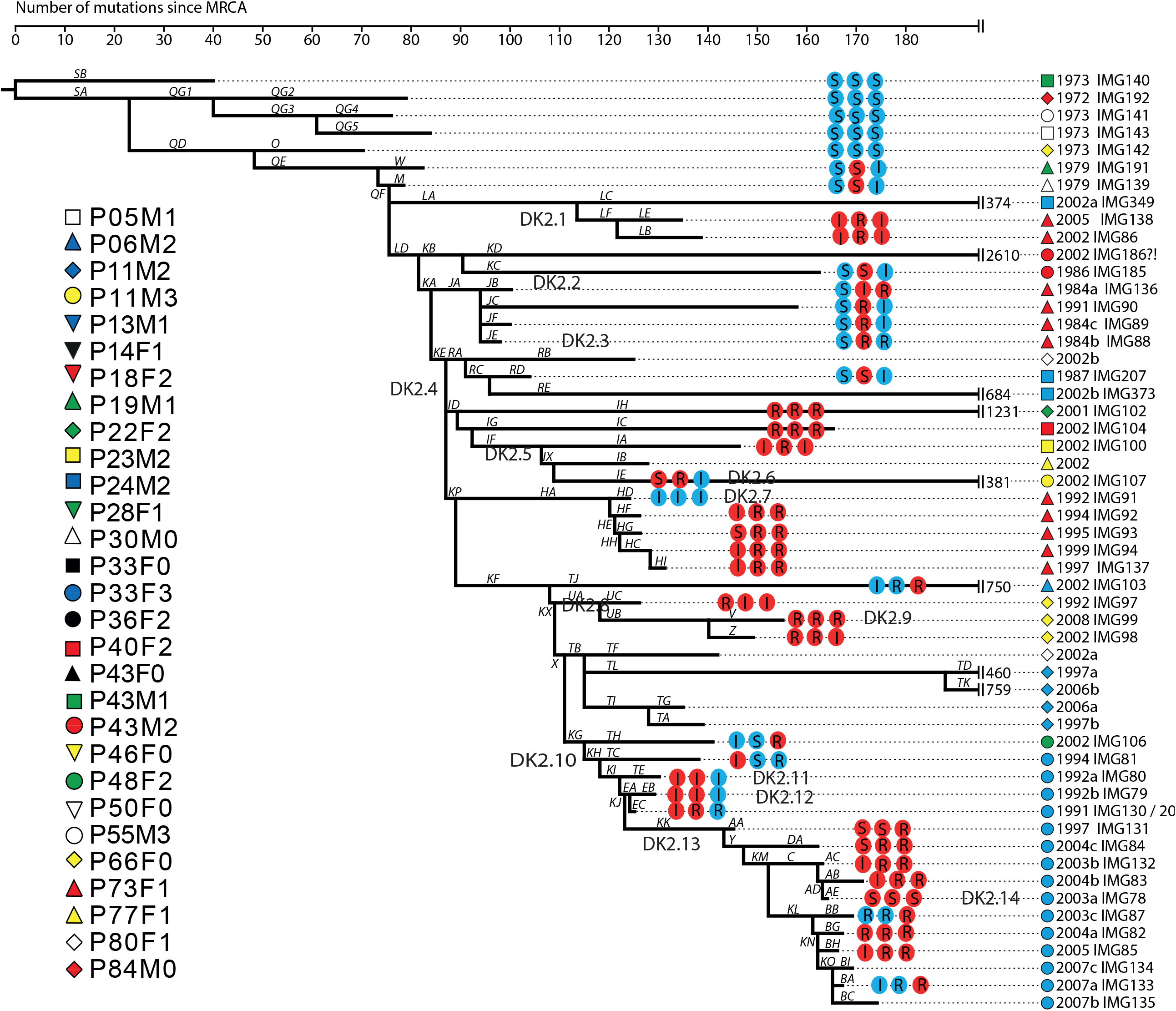

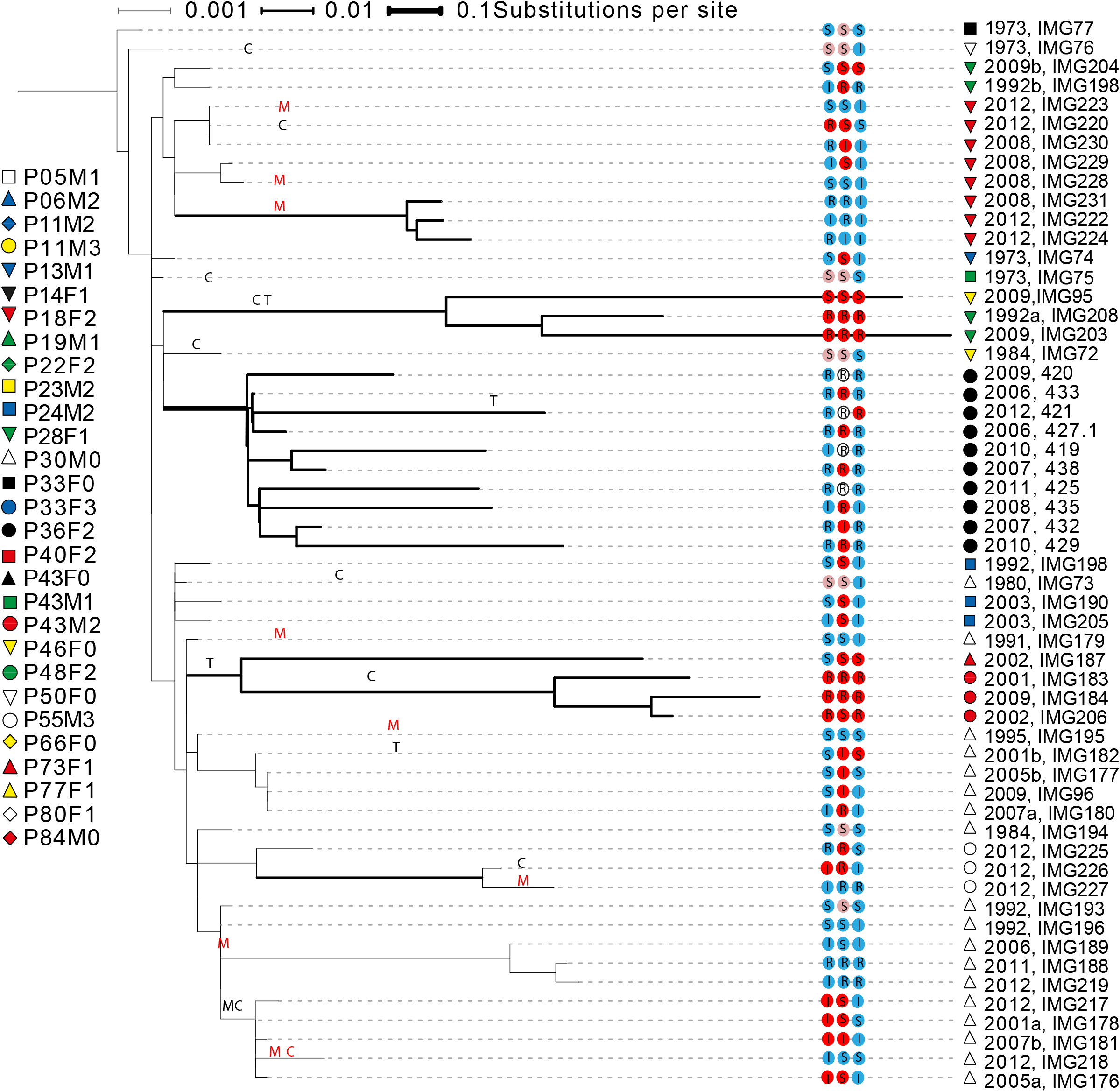

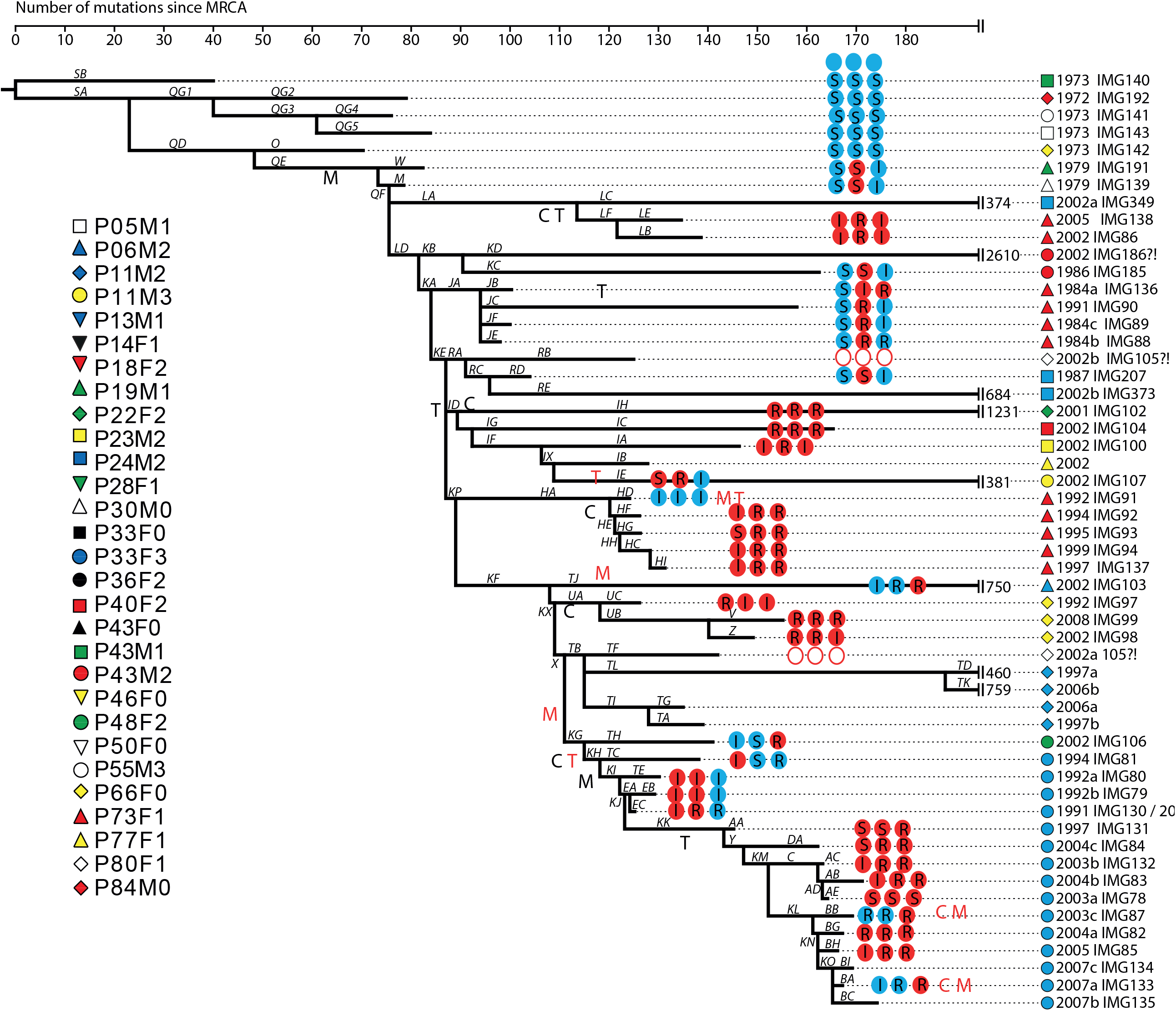

## References

1. Balaban NQ, Helaine S, Lewis K, Ackermann M, Aldridge B, Andersson DI, et al. Definitions and guidelines for research on antibiotic persistence (vol 17, pg 441, 2019). Nature Reviews Microbiology. 2019;17(7):460-.

2. Bigger JW. Treatment of staphylococcal infections with penicillin - By intermittent sterilisation. Lancet. 1944;2:497–500.

3. Hobby GL, Meyer K, Chaffee E. Observations on the mechanism of action of penicillin. P Soc Exp Biol Med. 1942;50(2):281–5.

4. Fisher RA, Gollan B, Helaine S. Persistent bacterial infections and persister cells. Nature Reviews Microbiology. 2017;15(8):453–64.

5. Bartell JA, Cameron DR, Mojsoska B, Haagensen JAJ, Pressler T, Sommer LM, et al. Bacterial persisters in long-term infection: Emergence and fitness in a complex host environment. Plos Pathogens. 2020;16(12).

6. Drescher SPM, Gallo SW, Ferreira PMA, Ferreira CAS, de Oliveira SD. Salmonella enterica persister cells form unstable small colony variants after in vitro exposure to ciprofloxacin. Sci Rep-Uk. 2019;9.

7. De Groote VN, Verstraeten N, Fauvart M, Kint CI, Verbeeck AM, Beullens S, et al. Novel persistence genes in Pseudomonas aeruginosa identified by high-throughput screening. Fems Microbiol Lett. 2009;297(1):73–9.

8. Viducic D, Murakami K, Amoh T, Ono T, Miyake Y. RpoN Promotes Pseudomonas aeruginosa Survival in the Presence of Tobramycin. Frontiers in Microbiology.2017;8.

9. Schumacher MA, Balani P, Min JK, Chinnam NB, Hansen S, Vulic M, et al. HipBA-promoter structures reveal the basis of heritable multidrug tolerance. Nature. 2015;524(7563):59–U108.

10. Isabella Santi PM, Enea Maffei, Adrian Egli, and Urs Jenal. Evolution of Antibiotic Tolerance Shapes Resistance Development in Chronic Pseudomonas aeruginosa Infections. mBio 2021;1(12):e03482–20.

11. Gollan B, Grabe G, Michaux C, Helaine S. Bacterial Persisters and Infection: Past, Present, and Progressing. Annu Rev Microbiol. 2019;73:359–85.

12. Balaban NQ, Merrin J, Chait R, Kowalik L, Leibler S. Bacterial persistence as a phenotypic switch. Science. 2004;305(5690):1622–5.

13. Olsen I. Biofilm-specific antibiotic tolerance and resistance. Eur J Clin Microbiol. 2015;34(5):877–86.

14. Orman MA, Brynildsen MP. Dormancy Is Not Necessary or Sufficient for Bacterial Persistence. Antimicrob Agents Ch. 2013;57(7):3230–9.

15. Levison ME, Levison JH. Pharmacokinetics and Pharmacodynamics of Antibacterial Agents. Infect Dis Clin N Am. 2009;23(4):791-+.

16. Van den Bergh B, Michiels JE, Wenseleers T, Windels E, Vanden Boer P, Kestemont D, et al. Frequency of antibiotic application drives rapid evolutionary adaptation of Escherichia coli persistence. Nature Microbiology. 2016;1(5).

17. Bakkeren E, Diard M, Hardt WD. Evolutionary causes and consequences of bacterial antibiotic persistence. Nature Reviews Microbiology. 2020;18(9):479–90.

18. Gardner A, West SA, Griffin AS. Is Bacterial Persistence a Social Trait? Plos One. 2007;2(8).

19. Kussell E, Leibler S. Phenotypic diversity, population growth, and information in fluctuating environments. Science. 2005;309(5743):2075–8.

20. Kussell E, Kishony R, Balaban NQ, Leibler S. Bacterial persistence: A model of survival in changing environments. Genetics. 2005;169(4):1807–14.

21. Levin-Reisman I, Gefen O, Balaban NQ. Tolerance and persistence promote the evolution of antibiotic resistance. Eur Biophys J Biophy. 2017;46:S73–S.

22. Levin-Reisman I, Ronin I, Gefen O, Braniss I, Shoresh N, Balaban NQ. Antibiotic tolerance facilitates the evolution of resistance. Science. 2017;355(6327):826–30.

23. Windels EM, Michiels JE, Fauvart M, Wenseleers T, Van den Bergh B, Michiels J. Bacterial persistence promotes the evolution of antibiotic resistance by increasing survival and mutation rates. Isme Journal. 2019;13(5):1239–51.

24. Bakkeren E, Huisman JS, Fattinger SA, Hausmann A, Furter M, Egli A, et al. Salmonella persisters promote the spread of antibiotic resistance plasmids in the gut. Nature. 2019;573(7773):276-+.

25. Mulcahy LR, Burns JL, Lory S, Lewis K. Emergence of Pseudomonas aeruginosa strains producing high levels of persister cells in patients with cystic fibrosis. J Bacteriol. 2010;192(23):6191–9.

26. Cohen NR, Lobritz MA, Collins JJ. Microbial Persistence and the Road to Drug Resistance. Cell Host & Microbe. 2013;13(6):632–42.

27. Marvig RL, Sommer LM, Molin S, Johansen HK. Convergent evolution and adaptation of Pseudomonas aeruginosa within patients with cystic fibrosis. Nature genetics. 2015;47(1):57–64.

28. Shrestha SD, Guttman DS, Perron GG. Draft Genome Sequences of 10 Environmental Pseudomonas aeruginosa Strains Isolated from Soils, Sediments, and Waters. Microbiol Resour Ann. 2017;5(34).

29. Jelsbak L, Johansen HK, Frost AL, Thogersen R, Thomsen LE, Ciofu O, et al. Molecular epidemiology and dynamics of Pseudomonas aeruginosa populations in lungs of cystic fibrosis patients. Infect Immun. 2007;75(5):2214–24.

30. Markussen T, Marvig RL, Gomez-Lozano M, Aanaes K, Burleigh AE, Hoiby N, et al. Environmental heterogeneity drives within-host diversification and evolution of Pseudomonas aeruginosa. MBio. 2014;5(5):e01592–14.

31. Marvig RL, Johansen HK, Molin S, Jelsbak L. Genome Analysis of a Transmissible Lineage of Pseudomonas aeruginosa Reveals Pathoadaptive Mutations and Distinct Evolutionary Paths of Hypermutators. Plos Genet. 2013;9(9).

32. Yang L, Jelsbak L, Molin S. Microbial ecology and adaptation in cystic fibrosis airways. Environ Microbiol. 2011;13(7):1682–9.

33. Rau MH, Marvig RL, Ehrlich GD, Molin S, Jelsbak L. Deletion and acquisition of genomic content during early stage adaptation of Pseudomonas aeruginosa to a human host environment. Environ Microbiol. 2012;14(8):2200–11.

34. Marley EF, Mohla C, Campos JM. Evaluation of E-Test for determination of antimicrobial MICs for Pseudomonas aeruginosa isolates from cystic fibrosis patients. J Clin Microbiol. 1995;33(12):3191–3.

35. Baker CN, Stocker SA, Culver DH, Thornsberry C. Comparison of the E-Test to Agar Dilution, Broth Microdilution, and Agar Diffusion Susceptibility Testing Techniques by Using a Special Challenge Set of Bacteria. J Clin Microbiol. 1991;29(3):533–8.

36. Robinson D. Convert Statistical Objects into Tidy Tibbles. https://broomtidymodelsorg/, https://githubcom/tidymodels/broomda. 2021.

37. Auguie B. gridExtra: Miscellaneous Functions for. “Grid” Graphics R package version 200. 2015.

38. Eklund A. The Bee Swarm Plot, an Alternative to Stripchart. https://githubcom/aroneklund/beeswarm. 2021.

39. Wickham H. Welcome to the Tidyverse. Journal of Open Source Software. 2019;4(43):1686.

40. Rossi E, La Rosa R, Bartell JA, Marvig RL, Haagensen JAJ, Sommer LM, et al. Pseudomonas aeruginosa adaptation and evolution in patients with cystic fibrosis. Nature Reviews Microbiology. 2020.

41. Hall CW, Mah TF. Molecular mechanisms of biofilm-based antibiotic resistance and tolerance in pathogenic bacteria. Fems Microbiology Reviews. 2017;41(3):276–301.

42. Andersen SB, Marvig RL, Molin S, Krogh Johansen H, Griffin AS. Long-term social dynamics drive loss of function in pathogenic bacteria. Proceedings of the National Academy of Sciences of the United States of America. 2015;112(34):10756–61.

43. Jorth P, Staudinger BJ, Wu X, Hisert KB, Hayden H, Garudathri J, et al. Regional Isolation Drives Diversification within Cystic Fibrosis Lungs. Pediatr Pulm. 2015;50:175–6.

44. Lopatkin AJ, Bening SC, Manson AL, Stokes JM, Kohanski MA, Badran AH, et al. Clinically relevant mutations in core metabolic genes confer antibiotic resistance. Science. 2021;371(6531):799-+.

45. Brazas MD, Breidenstein EBA, Overhage J, Hancock REW. Role of lon, an ATP-dependent protease homolog, in resistance of Pseudomonas aeruginosa to ciprofloxacin. Antimicrob Agents Ch. 2007;51(12):4276–83.

46. Goormaghtigh F, Van Melderen L. Single-cell imaging and characterization of Escherichia coli persister cells to ofloxacin in exponential cultures. Sci Adv. 2019;5(6).

47. Kohanski MA, Dwyer DJ, Collins JJ. How antibiotics kill bacteria: from targets to networks. Nature Reviews Microbiology. 2010;8(6):423–35.

48. Ciofu O, Beveridge TJ, Kadurugamuwa J, Walther-Rasmussen J, Hoiby N. Chromosomal beta-lactamase is packaged into membrane vesicles and secreted from Pseudomonas aeruginosa. J Antimicrob Chemother. 2000;45(1):9–13.

49. Medaney F, Dimitriu T, Ellis RJ, Raymond B. Live to cheat another day: bacterial dormancy facilitates the social exploitation of beta-lactamases. ISME J. 2016;10(3):778–87.

50. Bulitta JB, Ly NS, Landersdorfer CB, Wanigaratne NA, Velkov T, Yadav R, et al. Two Mechanisms of Killing of Pseudomonas aeruginosa by Tobramycin Assessed at Multiple Inocula via Mechanism-Based Modeling. Antimicrob Agents Ch. 2015;59(4):2315–27.

51. Van den Bergh B, Fauvart M, Michiels J. Formation, physiology, ecology, evolution and clinical importance of bacterial persisters. Fems Microbiology Reviews. 2017;41(3):219–51.

52. Pontes MH, Groisman EA. Slow growth determines nonheritable antibiotic resistance in Salmonella enterica. Sci Signal. 2019;12(592).

53. Lewis K. Persister cells, dormancy and infectious disease. Nature Reviews Microbiology. 2007;5(1):48–56.

54. Fridman O, Goldberg A, Ronin I, Shoresh N, Balaban NQ. Optimization of lag time underlies antibiotic tolerance in evolved bacterial populations. Nature. 2014;513(7518):418-0020+.

55. Stepanyan K, Wenseleers T, Duenez-Guzman EA, Muratori F, Van den Bergh B, Verstraeten N, et al. Fitness trade-offs explain low levels of persister cells in the opportunistic pathogen Pseudomonas aeruginosa. Molecular Ecology. 2015;24(7):1572–83.

56. Wong A, Rodrigue N, Kassen R. Genomics of Adaptation during Experimental Evolution of the Opportunistic Pathogen Pseudomonas aeruginosa. Plos Genet. 2012;8(9).

57. Martinez JL, Rojo F. Metabolic regulation of antibiotic resistance. Fems Microbiology Reviews. 2011;35(5):768–89.

58. Hansen E, Karslake J, Woods RJ, Read AF, Wood KB. Antibiotics can be used to contain drug-resistant bacteria by maintaining sufficiently large sensitive populations. Plos Biology. 2020;18(5).

59. Imamovic L, Sommer MOA. Use of Collateral Sensitivity Networks to Design Drug Cycling Protocols That Avoid Resistance Development. Sci Transl Med. 2013;5(204).

